# Transcriptomic view of key events during early embryogenesis in the duplicated Atlantic salmon genome

**DOI:** 10.64898/2026.08.03.742464

**Authors:** Prabin Sharma Humagain, Diego Perojil Morata, Maren Ileby Wessel, Jacob Seilo Torgersen, Sigbjørn Lien, Matthew Peter Kent, Daniel Macqueen, Victor Boyartchuk

## Abstract

Early embryogenesis is governed by tightly regulated transcriptional programs, including the maternal-to-zygotic transition (MZT), zygotic genome activation (ZGA), and maintenance of pluripotency. While these processes are well studied in model vertebrates, they remain poorly understood in salmonid fishes, whose genomes are shaped by a relatively recent whole genome duplication (WGD) event. Here, we present a temporally resolved transcriptomic analysis of Atlantic salmon (Salmo salar) embryogenesis using bulk RNA-seq across key stages spanning early embryogenesis. Dimensionality reduction and unsupervised clustering of gene expression revealed stage-specific transitions encompassing maternal RNA clearance, cell cycle regulation, and the onset of metabolic activity. We demonstrate that ZGA occurs early and in multiple phases, beginning soon after fertilization, accompanied by chromatin remodelling and the activation of epigenetic regulators. Duplicated gene pairs retained from the salmonid WGD frequently displayed asynchronous expression, indicative of functional divergence and the evolution of additional regulatory complexity of embryonic development. To gain insights into pluripotency, we integrated analyses of gene expression, transcription factor motifs, and chromatin accessibility, to reveal conserved regulators including genes encoding Pou5f3, Nanog, and Sox19b, alongside divergent functions of Klf family members. Our cross-stage profiling allowed us to define a novel panel of stably-expressed reference genes for normalization during quantitative PCR analyses, which were used to validate pluripotency- and differentiation-associated dynamics inferred by RNA-seq. Together, our findings delineate the transcriptional architecture of early embryogenesis in Atlantic salmon, revealing both conserved and lineage-specific features of pluripotency regulation, and providing a foundational resource for future functional genomics and stem cell applications in salmonids.

## Introduction

Recent advances in transcriptome profiling and omics have improved our understanding of embryonic development in model organisms (1–3), revealing conserved regulatory mechanisms and key molecular events (4,5). The first such critical event during early embryogenesis is the maternal-to-zygotic transition (MZT), defined as the shift in developmental control from maternally deposited transcripts to zygotic transcription and hence gene expression (6). A key component of MZT is zygotic genome activation (ZGA), which marks the onset of zygotic transcription and is a conserved feature of embryogenesis. The timing of ZGA, however, varies substantially across species (7). For instance, in mammals, ZGA can be seen 24 hours post-fertilization at the 2-4 cell stage. On the other hand, in *Xenopus*, *Drosophila*, and *Danio* models, ZGA is seen as early as 2-6 hrs, which corresponds to the 1024-4096 cell stages of development. Another important feature of early embryogenesis is the establishment and maintenance of pluripotency, enabling embryonic cells to differentiate into all somatic and germ cell lineages (8–10).

Despite some efforts to apply omic approaches in non-model species (11,12), our understanding of their embryonic transcriptome remains comparatively limited, particularly for species with life cycles that differ markedly from those of model organisms. Among such species is the Atlantic salmon (*Salmo salar*), a cold-adapted anadromous teleost of high ecological and economic importance (13,14). While extensive studies have focused on omic analysis of its later life stages, the molecular events defining early embryogenesis of Atlantic salmon remain poorly characterized (15). Understanding of pluripotency in Atlantic salmon likewise remains far behind that in mammals (16) and other model organisms.

Early development is sensitive to environmental conditions, particularly temperature (17,18). It can have lasting "carryover effects" that shape key traits throughout the life cycle of the organism, including its growth, survival, reproductive success, and overall fitness (19,20). As a cold-adapted species developing at temperatures well below those of conventional model organisms, Atlantic salmon may therefore follow distinct developmental trajectories that are not captured by existing frameworks. Therefore, benchmarking early development of the Atlantic Salmon at a high resolution is important to better understand normal development.

Atlantic salmon presents additional challenges in understanding early developmental processes due to its unique evolutionary history. The salmonid ancestor underwent a whole genome duplication (WGD) event approximately 88–103 million years ago, resulting in an autotetraploid genome (21). Although extensive rediploidization has occurred, many duplicated genes (ohnologs) remain, often exhibiting subfunctionalization or neofunctionalization (22). These duplicated gene pairs can have distinct expression patterns and regulatory roles during development, contributing further to the intricacy of early embryogenesis in this species. However, the impact of the salmonid WGD on embryonic transcriptome regulation remains poorly understood (23).

Systematic characterization of ZGA and pluripotency in Atlantic salmon does not only improve our understanding of its developmental biology but also provides new tools and markers for future applied research. In particular, insights into the regulation of pluripotency are foundational for the generation of induced Pluripotent Stem Cells (iPSCs)—somatic cells reprogrammed into a pluripotent state (24,25). iPSCs already have diverse applications, including functional genomics (26), disease modelling (27), regenerative therapies (28), and cell-based breeding platforms in aquaculture (29,30). However, to establish iPSC technology in Atlantic salmon, we still require additional of the underpinning mechanisms at play.

In this study, we use bulk RNA sequencing (RNA-seq) to investigate temporal transcriptomic dynamics of early embryogenesis in Atlantic salmon, while focusing on ZGA, pluripotency, and ohnolog expression patterns. By identifying key transcriptional shifts and regulatory gene sets, we elucidate the molecular architecture underlying normal embryonic development of this species. Overall, our findings fill a gap in salmonid developmental biology, laying the groundwork for novel iPSC and precision breeding applications.

## Methods

### Sample Collection and Embryo Processing

All Atlantic salmon (*Salmo salar*) embryos used in this study originated from the same commercial breeding stock provided by AquaGen AS. Embryos were generated and sampled at two institutions: the Norwegian University of Life Sciences (NMBU) and the Roslin Institute, University of Edinburgh (UoE). At UoE, embryos from a single developmental batch were collected across early embryonic stages as part of the AQUA-FAANG project. A subset of these samples was additionally processed and sequenced outside the AQUA-FAANG project (hereafter referred to as “UE samples”); however, both AQUA-FAANG and UE samples originated from the same developmental batch, genetic background, and incubation conditions. Publicly available AQUA-FAANG datasets therefore provide complementary stage coverage to the newly generated samples. Sample stages and origin are summarized in Figure 1.

**Figure 1.**
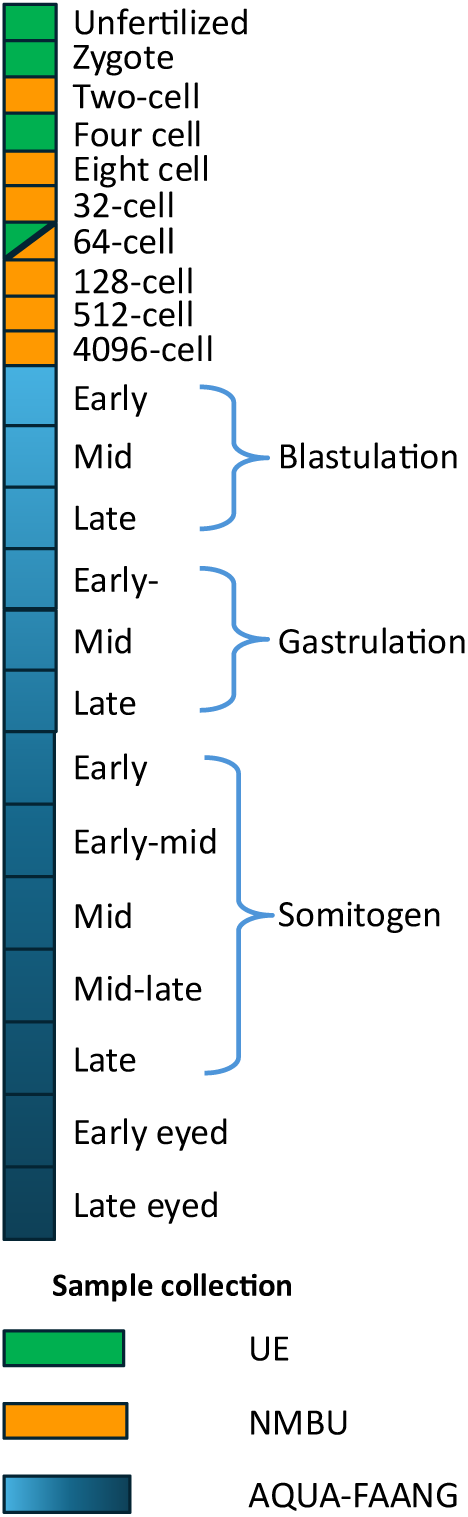
Figure showing the different developmental stages of Atlantic salmon used for analysis. The green and orange colour represents samples collected at the University of Edinburgh and NMBU. The blue colour represents the data supplemented from AQUA-FAANG project which were also collected at the same time as UE sample. Library preparation for the samples collected at NMBU were performed using non-stranded poly(A)-selected protocol.

Following fertilization, embryos were incubated at 4 °C (NMBU) or 8 °C (UoE) and monitored for developmental progression using brightfield microscopy. At NMBU, three biological replicate pools of 10 embryos were collected at defined developmental stages. Embryos were manually dechorionated in phosphate-buffered saline (PBS) under a dissecting microscope using fine forceps. Cells were carefully separated from the yolk sac and collected using a 100 µL pipette tip. Cell suspensions were pelleted by centrifugation at 100 × *g* for 5 min, the supernatant was removed, and cell pellets were stored at −20 °C for short-term storage prior to RNA extraction.

### RNA isolation and sequencing

Total RNA from NMBU samples was isolated using the RNeasy Mini Kit (Qiagen) according to the manufacturer’s instructions. RNA integrity was assessed using a TapeStation 4150 (Agilent), yielding RNA integrity numbers (RIN) between 8 and 10 for all samples. Libraries were prepared using a non-stranded poly(A)-selected protocol and sequenced by Novogene UK to a depth of approximately 30 million paired-end reads (PE150) per sample. The raw sequencing reads can be found under accession PRJEB98144 in European Nucleotide Archive (ENA) database.

For samples collected at the UoE (unfertilized, zygote, 4 cell and 64 cells – pool of 3 embryos grown in 8°C), total RNA was isolated using the TRIzol reagent (Qiagen) according to a standard operating protocol of the AQUA-FAANG project (23). Libraries were prepared for stranded paired-end sequencing and sequenced to a depth of ∼30 million PE150 reads per sample (Novogene UK).

In addition, publicly available RNA-seq datasets generated as part of the AQUA-FAANG project (PRJEB51855 in ENA database), spanning stages from early blastulation to late-eyed embryos, were downloaded and included in the analysis (see Supplementary Table 1 for details), which provide complementary coverage to our newly collected samples. All sequencing data—both newly generated and from AQUA-FAANG—were processed using the nf-core/rnaseq (v3.6) (31) of the *nf-core* collection of workflows, utilizing reproducible software environments from the Bioconda (32) and Biocontainers (33) projects. Transcript quantification and alignment were performed using the Salmo salar genome assembly Ssal_v3.1 (GCA_905237065.2), which includes 29 chromosomes and 4,222 unplaced scaffold regions. The corresponding Ensembl annotation contains 69,389 genes and 184,209 transcripts. Transcript alignment was performed using *STAR*(v2.6.1d) (34) and gene expression quantification was carried out using *salmon* (v1.5.2) (35). Read quality control was performed using *FastQC* (v0.11.9) (36), adapter and quality trimming was performed using *trimgalore* (v0.6.7) (37) and sorting and indexing of the alignments was carried out using *SAMtools* (v1.14) (38,39).

### Assessing the effects of different sample processing and library preparation

To assess for potential batch effects arising from different RNA extraction methods (column-based (RNeasy Mini Kit (NMBU)) vs. TRIzol-based (UoE)) and library preparation protocols (stranded vs. unstranded), we performed pairwise differential expression (DE) analyses between the available developmental stages, comparing each sampled stage with the next in the developmental sequence (e.g. zygote vs. two-cell, two-cell vs. four-cell, etc). DE analysis was conducted using *DESeq2* (v1.40.2) (40) on raw transcript counts. Genes with a log2 fold change (LFC) > 1.0 and adjusted *p* < 0.05 were considered significantly differentially expressed.

To identify genes likely affected by technical variation rather than true biological differences, we examined DE results across comparisons of adjacent developmental stages that had been sampled and processed using different extraction or library preparation methods. We then identified a subset of differentially expressed genes that were common to all 6 contrasts generated using different library preparation methods. Because of their persistence across stages, we assumed that these DE genes are artifacts attributed to differences in methodology. We therefore excluded them from downstream analyses.

Additionally, because unstranded RNA-seq protocols are known to misattribute expression levels in regions where transcripts overlap on opposite strands (41), we flagged excluded genes that overlapped with antisense transcripts as potentially affected by this issue.

### Identification of persistent and new gene expression

As early embryogenesis is characterized by precise regulation of gene expression, our goal was to capture the dynamic transcriptional changes in genes throughout development and track the emergence of novel transcripts. To achieve this, we used Transcripts Per Million (TPM) values of each gene as the measure of gene expression. However, while calculating TPM, spike-in normalization was not employed. For each developmental stage, the mean TPM per gene was calculated across three biological replicates (n = 3) for those genes which had TPM>0.5 for at least 2 replicates. A gene was considered expressed at a specific stage if its average TPM exceeded 2; this threshold was chosen to minimize the influence of low-level transcriptional noise while capturing biologically relevant expression. We applied an additional filter to define a set of persistently expressed genes. Specifically, a gene was retained only if it met the TPM > 2 threshold in three consecutive developmental stages. This criterion was chosen to exclude spurious transcripts and to focus the analysis on genes with sustained transcriptional activity. The resulting list of persistently expressed genes formed the basis for all downstream expression analyses. In the meantime, the list of genes that were excluded were clustered using Self Organizing Map (SOM) using the *kohonen* package (Version 3.0.12) (42) clustering to identify the developmental stages where their expression peaked.

Using the curated list of persistently expressed genes, we identified newly expressed genes for each developmental stage. A gene was defined as newly expressed at a given stage if it was not expressed (TPM ≤ 2) in all prior stages and exceeded the TPM > 2 threshold at the current stage. To assess the potential biological relevance of these newly expressed genes at their specific stages of expression initiation, we performed GO analysis.

### GO analysis

Gene Ontology (GO) enrichment analysis was performed using the *enrichGO()* function from the *clusterProfiler* Bioconductor package (v4.12.0) (43). Enrichment was carried out separately for Biological Process (BP) and Molecular Function (MF) categories. For each gene set of interest, enrichment was tested using a hypergeometric test against a background universe of all genes expressed across the dataset. GO annotations were derived from a custom organism-specific annotation package (org.Ssalar.eg.db) for *Salmo salar*, built using AnnotationForge (v1.46.0) (44) with Ensembl gene IDs as the key.

After enrichment, results were filtered to retain only GO terms with adjusted p-value < 0.05 (Benjamini–Hochberg method) and fewer than 400 annotated background genes to remove overly broad terms. Redundant GO terms were further reduced using two complementary strategies:

First, semantic similarity-based reduction using the *rrvgo* package (45), applying the "Rel" similarity metric with a threshold of 0.7 to retain the most representative terms followed by Gene content-based clustering, where GO terms sharing significant overlap in gene annotations (Jaccard index) were hierarchically clustered (cut height adjusted for different GO analysis), and only the most significant GO terms per cluster was retained. The final set of non-redundant GO terms was visualized using dot plots, with point size representing gene count and colour encoding statistical significance (–log10 adjusted *p-*value). Plots were generated using *ggplot2* (v3.5.1) (46).

### Chromosomal Enrichment of Newly Expressed Genes

For each developmental stage, a chi-square goodness-of-fit test was used to assess whether newly expressed genes were distributed across chromosomes in proportion to the total gene content of each chromosome. Chromosomes 1–29 were analyzed individually, while all non-chromosomal scaffolds were combined into a single category to ensure adequate expected counts. For each stage, chromosome-specific deviations were evaluated using standardized residuals derived from the chi-square test. Two-sided p-values were calculated from the standardized residuals using the normal approximation and adjusted for multiple testing across chromosomes using the Benjamini–Hochberg procedure. Chi-square p-values were additionally adjusted across developmental stages using the Benjamini– Hochberg method.

### Characterization of Ohnolog Expression Dynamics during embryogenesis

Given the Atlantic salmon’s evolutionary history of WGD, identifying duplicated gene pairs (ohnologs), to explore whether newly expressed genes included these ohnologs, and whether their expression diverged, we performed a comparative genomic analysis between Atlantic salmon and a phylogenetically close sister species, Rainbow trout (*Oncorhynchus mykiss)*. For this, we obtained a curated orthology table between Atlantic salmon and Rainbow trout from SalmoBase (23). Then, we focused on ohnolog pairs present as exactly two copies in both species to minimize ambiguity in duplication origin. To ensure evolutionary conservation and biological relevance, only those gene pairs that also exhibited synteny— defined as conserved gene order across species—were retained. This filtering yielded a high-confidence list of 21,042 syntenic ohnolog genes for further analysis.

We limited our analysis to focus on ohnologs that were expressed in stages up to, and including, early blastulation, and noted the developmental stage at which each gene in the ohnolog pair first became transcriptionally active.

Based on their expression onset, ohnolog pairs were classified into the following categories: (i) synchronous activation, in which both genes initiated expression at the same developmental stage; and (ii) asynchronous activation, encompassing cases a - where both genes were newly expressed before blastulation but at different stages, b - where one gene was maternally expressed and the other zygotically expressed, c - where one gene initiated expression before and the other after blastulation, or d - where only one gene of the pair was expressed during embryogenesis.

To test whether ohnolog genes were over-represented among newly expressed genes, we compared the observed proportion of ohnolog genes within the set of newly expressed genes to their genome-wide frequency using a one-sided binomial test. This test assessed whether newly expressed genes were more likely to be retained duplicates than expected by chance.

To evaluate divergence in expression timing between ohnolog pairs, we focused on pairs in which both duplicates initiated expression prior to early blastulation. The number of synchronously initiating pairs was compared to expectations under a simplified null model assuming equal probability of expression initiation across developmental stages using a one-sided binomial test. This analysis tested whether synchronous initiation occurred more frequently than expected under random stage assignment.

### Visualization of the closeness of biological replicates across stages

To visualize the data dispersion of our samples, we performed a Principal Component Analysis (PCA) using the raw counts with R package ‘stats’ (version 4.4.0) and plotted the data using *plot3d* function from the R package *rgl* (version 1.3.1) (47). Based on the loading values for the first three Principal Components (PCs) -PC1, PC2 and PC3, the 200 genes with the highest loading value (both positive and negative) for respective principal components were analysed for their enriched biological and molecular function, again using the *clusterProfiler* package for GO term analysis.

### Clustering of the genes based on their expression profile across the early developmental stages

To explore expression dynamics during development, persistently expressed genes were clustered based on their expression pattern using SOM clustering. Each SOM cluster was analysed using the *clusterProfiler* package to identify enrichment of Gene Ontology terms.

### Identification of pluripotency genes

To identify pluripotency genes, we first applied a candidate gene approach. For this, a list of stem cell markers was compiled from the literature that defined pluripotency genes in other organisms including human, mouse (48)(49) and teleost models (50). The corresponding orthologues in the Atlantic salmon were looked up using gene names assigned to the orthologues in Atlantic salmon, Zebrafish, Rainbow trout and human in descending order of priority. The orthologue list for the three lookup organisms was obtained using Ensembl *biomaRt* tool (51,52).

Candidate genes were further expanded using a *de novo* approach to identify unannotated Atlantic salmon pluripotency genes. For this, two subsets of developmental stages were defined, one that is likely to predominantly contain pluripotent cells (4,096 cell stage to late blastula) and the other where differentiation has begun (late gastrulation to late somitogenesis stage embryo). Using the Bioconductor *DESeq2* package, we performed a DE analysis between the pluripotency and differentiated subsets of embryo samples. The differentially regulated genes were defined as those with |LFC|>1 and adjusted p-value < 0.05.

### Identification of transcription factor binding motifs

To identify Atlantic salmon pluripotency regulators, we focused on POU domain-containing transcription factor (Pou5f3, also called OCT4 or Pou5f1), a well-defined master regulator of pluripotency across mammals and fish (53,54). We described the promoter region as the sequence ranging from 2500 bp upstream of the transcription start site (TSS) to 1000 bp downstream of TSS. Since the TSS can be different based on the isoforms, we focused only on the longest transcript. We got the position weight matrix (PWM) file for the motifs from the website https://jaspar.elixir.no/ for the *pou5f3* gene (MA1115.2) and a composite motif for Pou5f3:Sox2 (SRY-box 2) (MA0142.1) corresponding to (5’-ATGCAAAT-3’) and (Sox binding motif: 5’-CATTGTT-3’) respectively. The compositive motif of Pou5f3: Sox2 was also used because the literature shows that Pou5f3 functions with Sox2 among other factors to control the expression of different genes (55). SRY-box 2 (Sox2) is a transcription factor and core pluripotency maintenance gene in mammals, however, in zebrafish, another Sox family member (Sox19b) was reported to perform the same function (8). We searched for the occurrence of these motifs in the promoter regions of all the genes using matchPWN package (version 2.40.2) (56) with min. match = 90% as criteria. To limit our search, we filtered for only the motifs based on the specific chromatin state, as described in https://salmobase.org/datafiles/datasets/Aqua-Faang/robust_ATAC_peaks/unified_annotated_peaks/AtlanticSalmon_unified_peaks.bed (2024-05-30), ATAC-seq data from the late blastulation stage, from the AQUA-FAANG project (57). Only the genes which have the TF binding motifs in the regions – ‘Accessible chromatin’, ‘Active TSS’, ‘flanking upstream’ or ‘Active enhancer’ were included. These genes were then cross-referenced with the genes upregulated in the pluripotent stages of the Atlantic salmon development to identify those potentially regulated by Pou5f3 alone or by Pou5f3: SoxB1 family combined.

### Validation of observed gene expression

For validation of the expression patterns observed in candidate pluripotency and differentiation genes, quantitative polymerase chain reaction (qPCR) was used. To enable accurate normalization, novel housekeeping genes (HKGs) were identified from our transcriptomic dataset based on comparatively high mean TPMs and low variance during embryogenesis.

A new set of validation samples was collected while keeping the eggs/embryos at 4ᵒC, as this was done at Cigene. The samples were collected at 2 different time points: late-blastulation and early/mid-gastrulation (n=3 biological replicates). Ten embryos were collected for each replicate and total RNA was collected using the RNeasy micro kit (Qiagen, 74104) following the manufacturer’s instruction and as described above for RNA-Seq.

Total RNA concentation was measured using a NanoDrop 8000 (Thermo Fisher Scientific) with 800 ng used for cDNA synthesis using iScript™ cDNA Synthesis Kit (Cat no.: 1708890; Bio-Rad) following the manufacturer’s instruction in 20µl reactions for each sample. 1 µl of cDNA was used as template in qPCR using SsoAdvanced™ Universal SYBR® Green Supermix (Cat.no.: 1725271; Bio-Rad). The names and sequences of qPCR primers for quantification of selected pluripotency, differentiation and HKGs are provided in Table 1. The expression of other genes tested was normalized to house-keeping genes and compared between samples.

**Table 1.**
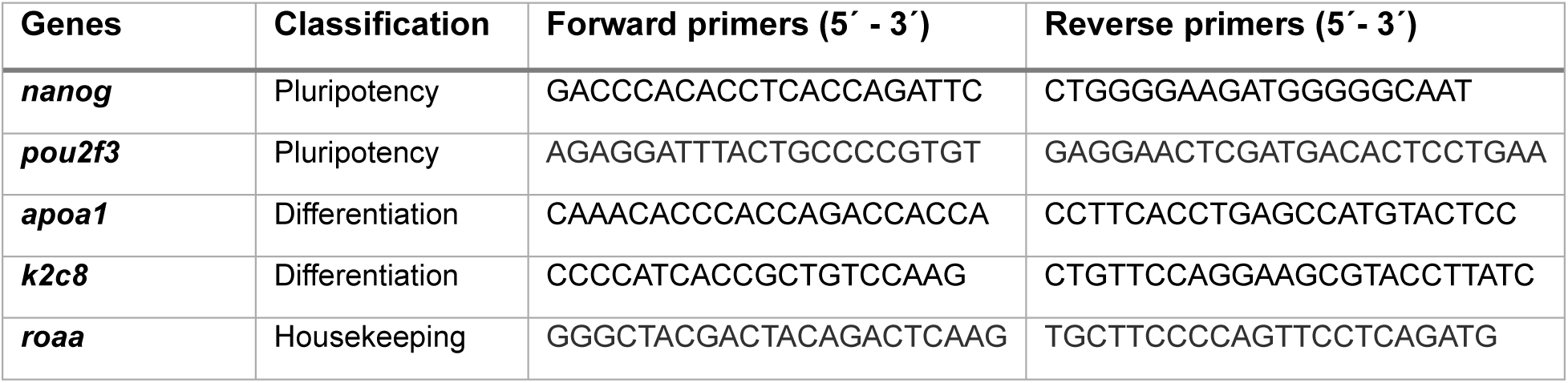
Primers used for quantitative PCR (qPCR) of candidate pluripotency and differentiation marker genes, along with housekeeping genes.

## Results

### Sequencing data across methodologies shows high consistency

To obtain a comprehensive map of embryonic gene expression in Atlantic salmon, we compiled an RNA-seq dataset from discrete stages and supplemented it with publicly available RNA-seq data (Supplementary Table 1). The combined dataset included samples generated under different rearing temperatures (4°C and 8°C) and using distinct library preparation methods.

An earlier study by Zhao et al. reported that choice of the sequencing method impacts analysis of differential gene expression (41), with strand-specific information enhancing gene annotation, particularly for overlapping genes. To test the presence of such effects, we quantified gene expression across the potentially affected datasets using an identical RNA-seq pipeline (see Methods). Subsequently, we performed pairwise differential expression (DE) analysis between the adjacent developmental stages. The results of the DE analysis are summarized in Figure 2.

**Figure 2.**
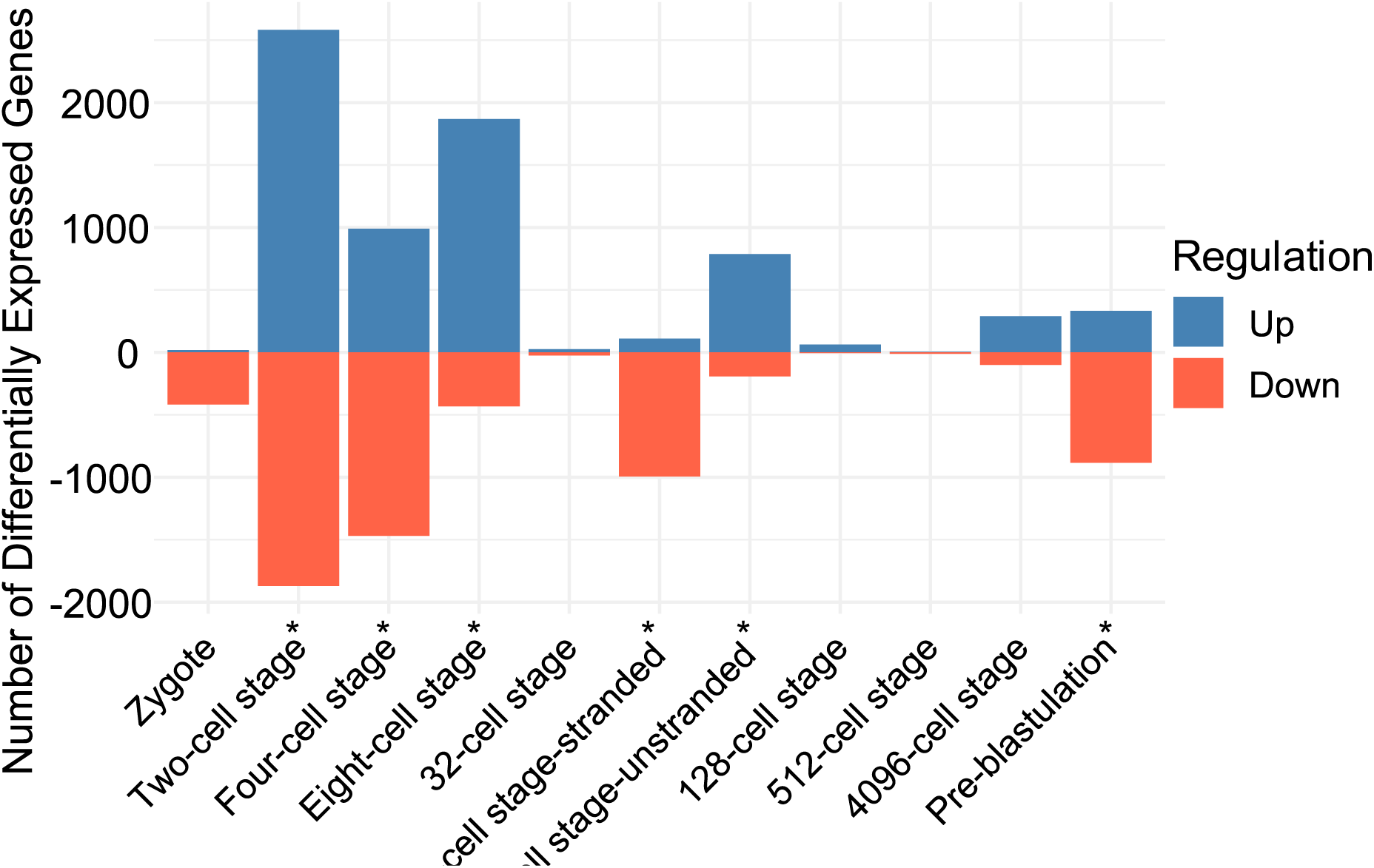
Summary of the number differentially expressed genes in different developmental stages of Atlantic salmon embryogenesis when compared to the preceding developmental stage. *Developmental stages where DE analysis was between the samples from stranded- and unstranded-library preparation and were used to identify the effect of different library preparation method – common differentially expressed genes.

Among the DE genes between the adjacent developmental stages, we found 489 genes that were consistently varying between samples that used different methodologies to collect the data. Of these, 477 overlapped with other annotated genes, including 429 overlapping genes genes transcribed from the opposite strand. This observation is likely due to artifacts introduced during alignment of sequencing reads rather than true biological differences and were excluded from further analysis. Because only a minority of overlapping genes were differentially quantified in datasets that used different methodology, we concluded that we could combine theses datasets for our analysis.

### Principal Component Analysis (PCA) of Gene Expression

As a first step towards characterising the early embryonic transcriptome of Atlantic salmon, we filtered our data to identify persistently expressed genes by averaging transcript abundance (TPM) values from pooled biological replicates (only genes with TPM>0.5 in at least 2 replicates) and excluding genes with average TPM ≤ 2. This yielded 31,746 expressed genes overall in the dataset. To reduce noise, only genes expressed across at least three consecutive stages were classified as “persistently expressed” which resulted in a final set of 29,150 genes.

To visualize the sample scatter across studied developmental stages, we performed PCA. The first three PCs explained 58.7% of the observed variance in gene expression (Figure 3). The biological replicates clustered closely, and samples showed a clear progression corresponding to developmental time.

**Figure 3.**
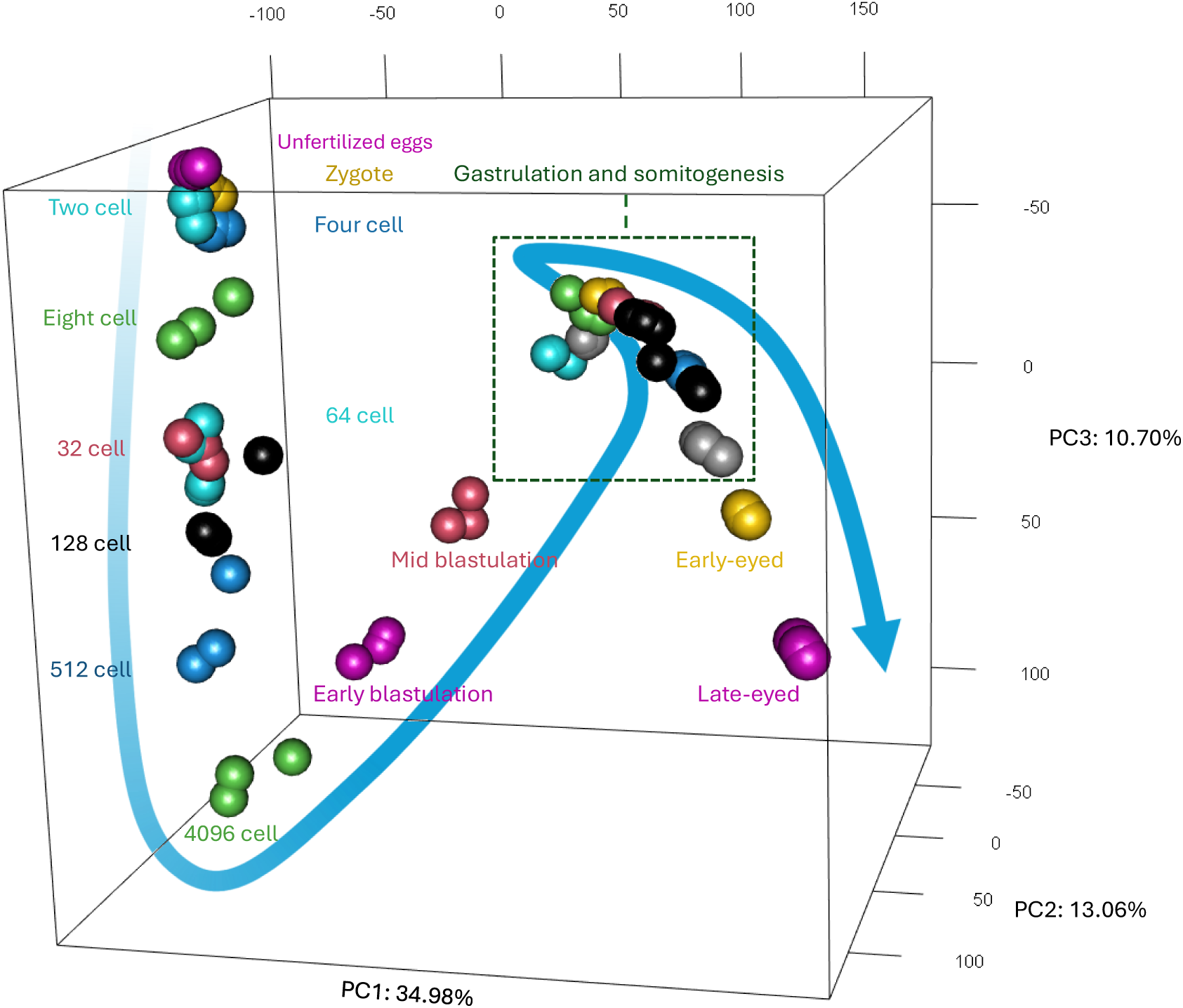
3D PCA of the complete dataset. Blue ribbon follows the developmental progression

To interpret biological contributions to each PC, we performed GO enrichment analysis with the 2.5% genes with the highest absolute loading scores for PC1, PC2, and PC3 (Supplementary Figure 1, Supplementary table 2).

### Stage specific expression of genes

A total of 2,596 genes were expressed for less than 3 consecutive stages and were excluded from the persistently expressed gene set (Supplementary Table 3). Clustering of these excluded genes showed a stage specific expression for majority of them (Supplementary Figure 2). Genes associated with the complement system were enriched in cluster 1 (unfertilized eggs). Genes involved in RNA splicing and rRNA modification, including members of the SNORA family, were predominantly expressed during early blastulation stages (cluster 4), while spliceosomal RNAs were detected primarily during the pre-blastulation stage (cluster 3). Though most of these excluded genes were expressed at low abundance, 107 showed TPM values greater than 10 in at least one developmental stage.

### Identification of newly expressed genes and Zygotic Genome Activation (ZGA)

To define the timing of ZGA in Atlantic salmon, we identified the earliest developmental stage at which transcripts absent from earlier stages, including unfertilized eggs, were stably expressed. First, the number of expressed genes was quantified at each developmental stage (Figure 4 and Supplementary Table 4). The unfertilized egg contained transcripts mapped to 15,376 genes, which decreased to 14,632 in the zygote and 14,312 at the 2-cell stage. From this point onward, a progressive increase in gene expression was observed, peaking at the late-eyed stage (24,631 genes). Approximately, 70% of genes expressed in each stage were annotated in ensemble database.

**Figure 4.**
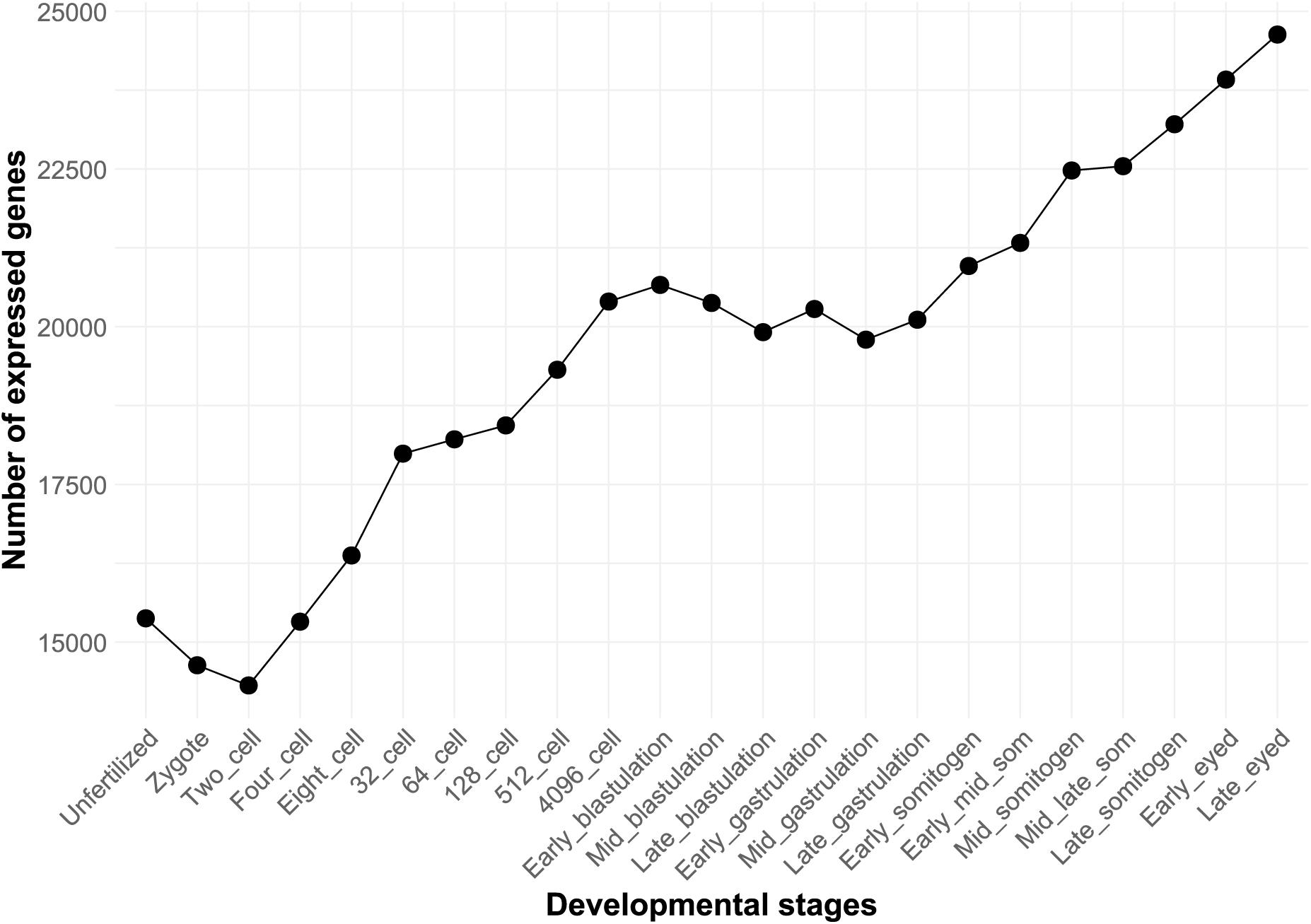
The total number of expressed genes (persistently expressed in at least 3 consecutive stages) at each developmental stage in Atlantic salmon embryogenesis (TPM>2).

Comparison of successive developmental stages revealed a progressive emergence of new transcripts mirroring the increasing complexity of embryo. Despite an overall decrease in the total number of genes scored as present from an unfertilized egg (15,376) to zygote (14,632), we found 143 newly expressed genes in the zygote. The number of newly expressed transcripts steadily increased in the next two stages peaking at the 8-cell stage (1,413) (Figure 5). This observation strongly suggests that in Atlantic salmon ZGA occurs very early in development with a substantial number of zygotically expressed transcripts detected already at the two-cell stage.

**Figure 5.**
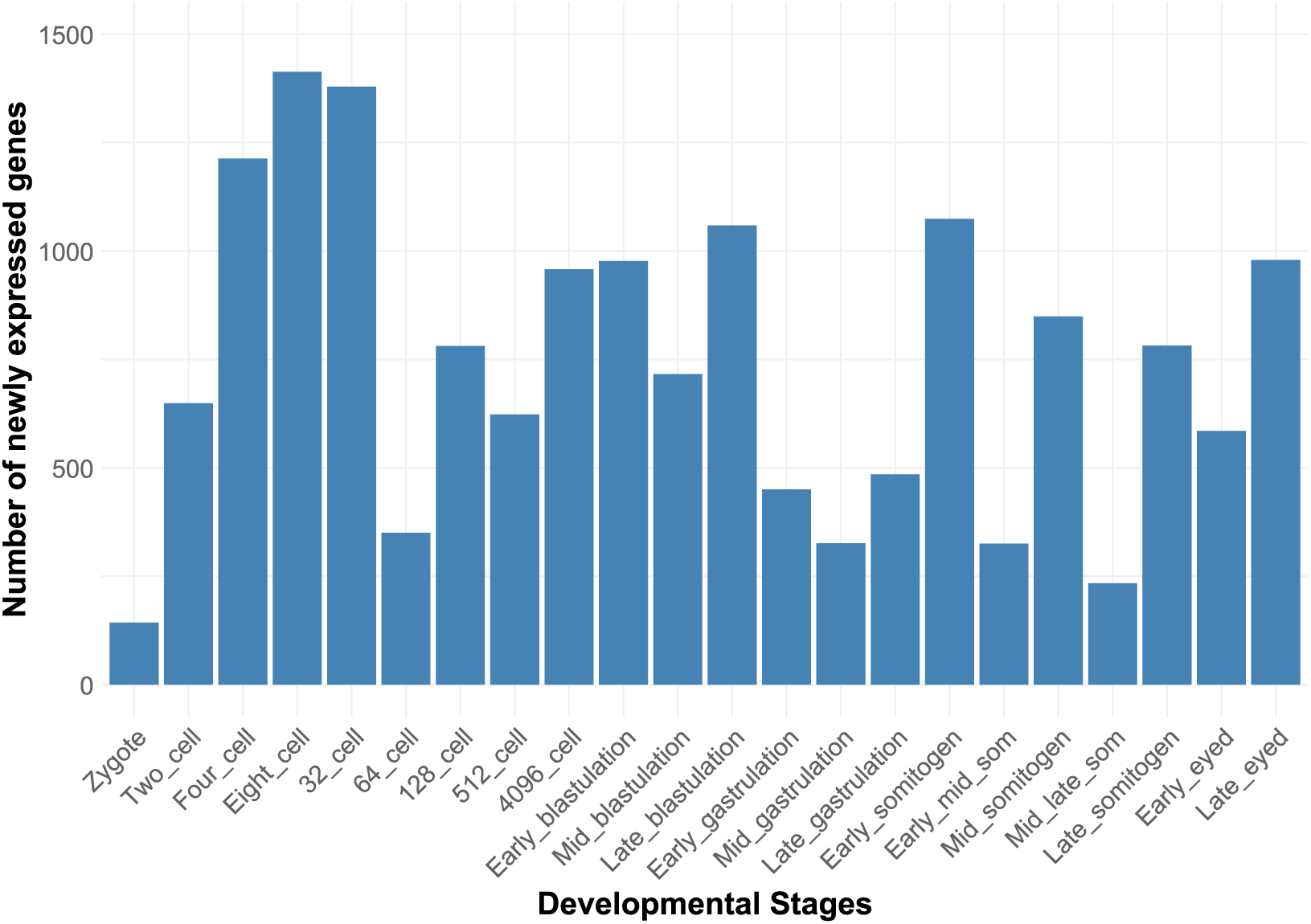
Inferred dynamics of zygotic gene expression initiation based on the first appearance of DEGs compared to the prior developmental stage. Only genes that are expressed persistently for three consecutive stages were considered.

### Enrichment Analysis of Newly Expressed Genes

To assess whether genes that are newly expressed at each developmental stage share common functions, we performed GO enrichment analysis for Biological Process and Molecular Function terms. We found that stage-specific gene sets exhibited distinct functional enrichments. Consistent with requirements for initiation of zygotic transcription, chromatin remodelling molecular functions were overrepresented in transcripts, including those for *chd8*, *smarca1* and *smarca2*, that appeared at the two-cell stage (Figure 6B). At the 8-cell stage, newly expressed genes were enriched for GTPase-related protein kinase activity. The 128-cell stage marked emergence of genes enriched for GO terms related to tRNA processing and DNA recombination functions. Full enrichment results are provided in Supplementary Table 5.

**Figure 6.**
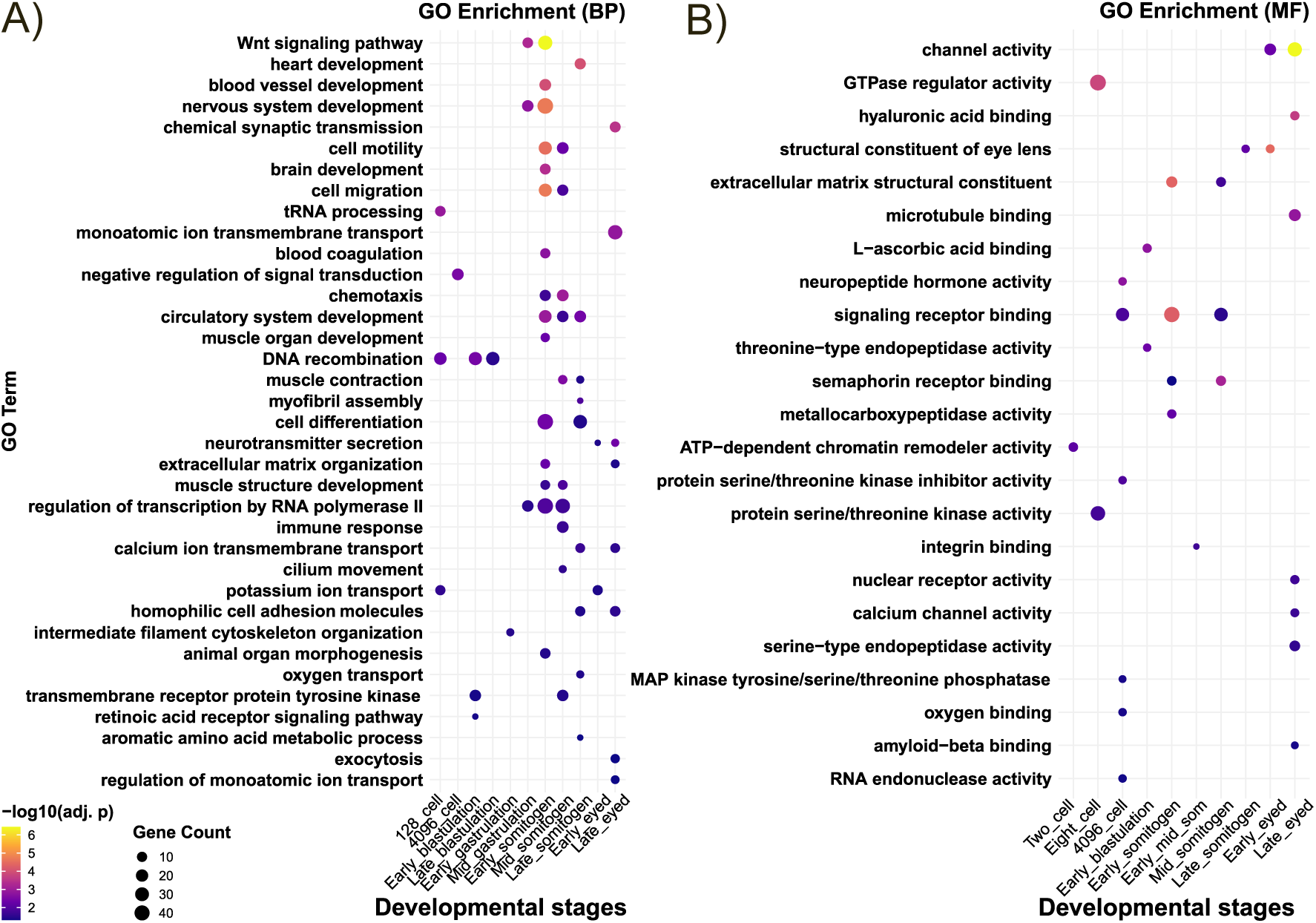
Gene ontology analysis of newly expressed genes for stages with significant enrichment of GO terms during early embryogenesis. A) Enrichment of biological processes. B) Enrichment of molecular functions. *p-value can only be compared withing the stage and not between the stages.

As previous studies have identified chromosomal clustering of a MZT-associated gene block in zebrafish (58), we examined whether newly expressed genes in Atlantic salmon were non-randomly distributed across chromosomes. For this, we mapped newly expressed genes to their genomic location and assess the chromosomal enrichment. This analysis revealed a non-uniform distribution of newly expressed genes across the genome, with several chromosomes showing significant under- or over-representation (chi-squared test). Notably, chromosome 2 was consistently overrepresented across multiple early developmental stages (Figure 7; Supplementary Table 6).

**Figure 7.**
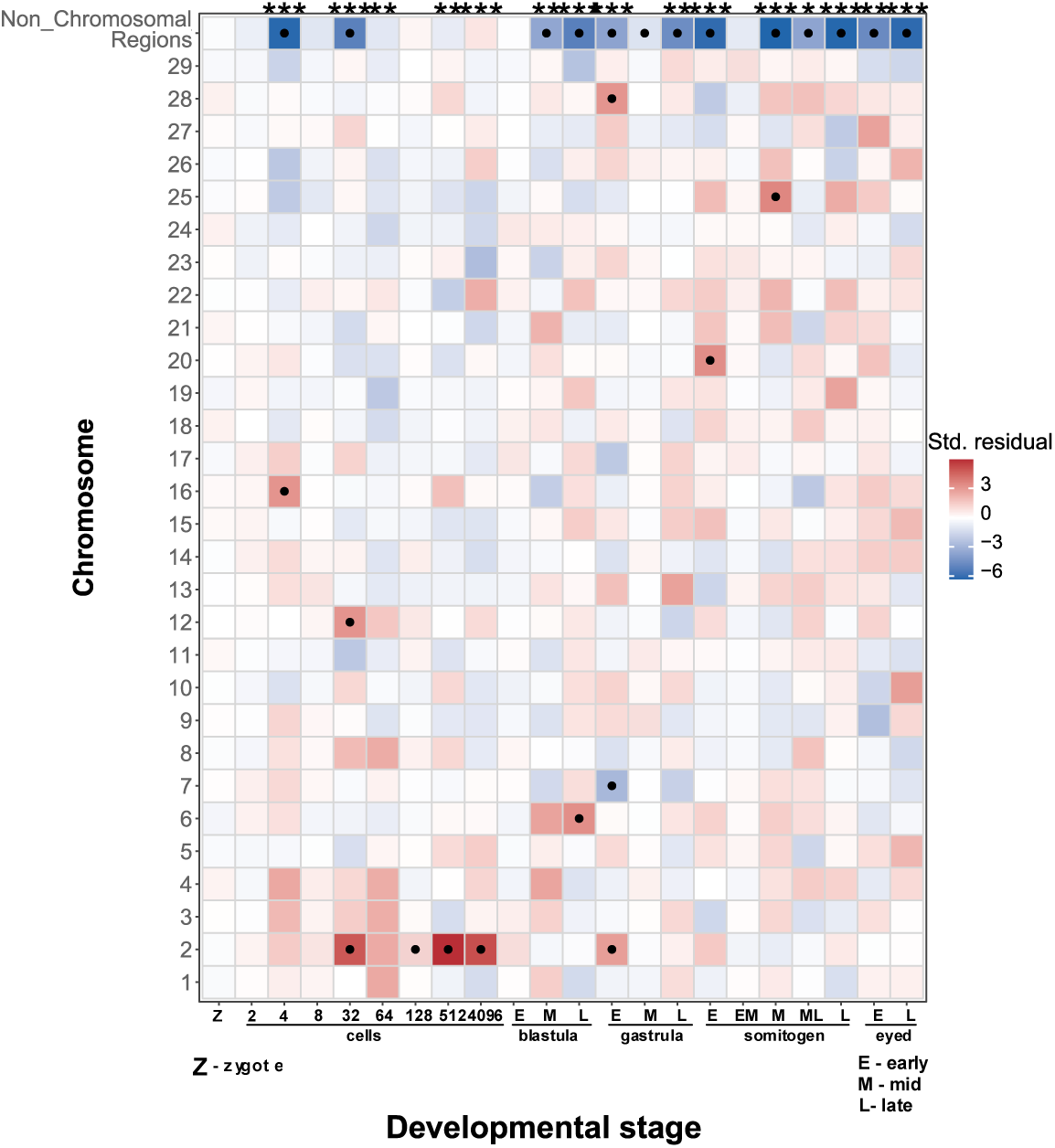
Chromosome-level enrichment of newly expressed genes across early embryonic stages. Heatmap shows standardized Pearson residuals from chi-square tests comparing the observed number of newly expressed genes per chromosome to the expected number based on chromosome-wide gene content at each developmental stage. Positive residuals (red) indicate over-representation, while negative residuals (blue) indicate under-representation of newly expressed genes on a given chromosome. Black dots denote chromosomes with statistically significant deviations from expectation after Benjamini–Hochberg multiple-testing correction (adjusted p < 0.05) within each stage. Developmental stages with a significant overall chromosome-level distribution (chi-square test, Benjamini–Hochberg adjusted p < 0.05) are indicated by stars above the heatmap (p < 0.05; p < 0.01; p < 0.001). All non-chromosomal scaffolds were grouped into a single category (“Non-Chromosomal Regions”).

### Ohnologs showing asynchronous activation

Many ohnolog pairs retained from the salmonid WGD exhibit divergent expression patterns, even within the same cell type (59,60). We therefore examined whether ohnolog duplicates show divergence in the timing of expression initiation during early embryogenesis. Our goal was to identify patterns of divergent expression between duplicated genes and to gain insights into possible functional divergence of ohnolog post-WGD during early development.

Of 8,486 genes that initiated expression up to early blastulation, a developmental transition marking the onset of widespread zygotic transcription, 4,097 belonged to annotated ohnolog pairs. This represents a significant overrepresentation relative to the genomic background (binomial test, p = 5.78e-262) and suggests that newly expressed genes are predominantly retained duplicates.

Among 802 ohnolog pairs, for which both copies of duplicates initiated expression prior to blastulation, only 209 pairs were synchronously expressed within the same developmental stage. This number is significantly lower than expected under a null model assuming equal initiation probability across stages (binomial test, p = 5.5e-32). In contrast, 593 pairs initiated expression asynchronously prior to blastulation, and an additional 678 pairs exhibited split initiation, with one duplicate initiating expression before and the other after blastulation.

Finally, 1,322 maternally deposited genes had ohnolog partners that initiated expression by early blastulation, further highlighting extensive decoupling of expression timing between duplicated genes (Supplementary Figure 3).

Our analysis defined 7 classes of ohnologs transcriptional activation. By analyzing genes in each separate class for enrichment of GO terms, we found functional differences between the maternal/zygotic copies of ohnolog pair as well as the split group of before-/after blastulation copies (Supplementary Figure 4). This observed difference in the enrichment suggests that there is divergence in ohnolog pair function during development.

### Clustering of RNA-seq data showed distinct groups of gene expression patterns

To obtain an overview of temporal gene expression patterns, we analyzed 12 distinct gene sets defined by SOM clustering of scaled transcript abundance data (Figure 8). GO term enrichment analysis of these clusters provided novel insights in timing of initiation of transcription of genes supporting essential developmental functions (Supplementary Table 7 including genes in each cluster, summarized in Supplementary Figure 5).

**Figure 8.**
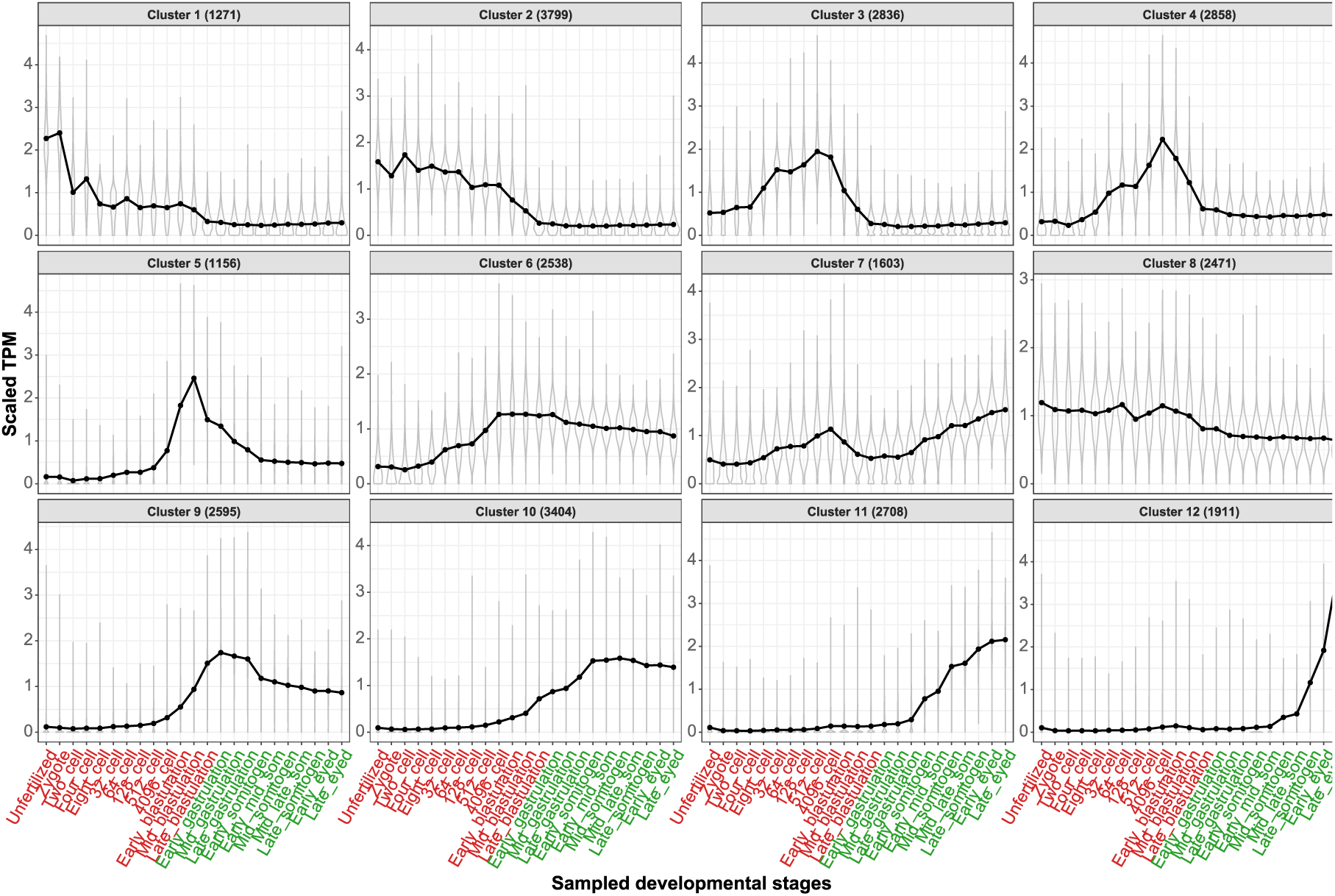
Clustering of all the genes based on their expression profile. Y-axis shows the scaled (but not centered zero) values for expression. Number of genes in each cluster are provided inside parenthesis.

Clusters 1 (1,274 genes) and 2 (3,799 genes) were dominated by maternally deposited transcripts, showing high abundance in unfertilized eggs followed by progressive degradation, reaching minimal levels by late blastulation. These genes were enriched for functions related to cell cycle regulation, chromosomal organization, DNA polymerase activity, intracellular protein transport, and fatty acid metabolic processes.

In contrast to clusters composed of maternal transcripts, clusters 3 (2,836 genes), 4 (2,858 genes), and 6 (2,538 genes) showed increasing expression early in development. Genes in these clusters were enriched for terms encompassing processes of protein ubiquitination, DNA repair, RNA processing, ribosome biogenesis, and phosphatidylinositol metabolism, which is consistent with expected requirement of early zygotic development, A similar but delayed expression profile is seen in cluster 5 (1,156 genes) genes, with a sharp increase in expression beginning at the 64-cell stage and peaking at mid-blastulation, suggesting a preparatory role in oncoming differentiation. This set of genes were enriched in molecular functions relating to protein kinase inhibitor and endopeptidase activity. In contrast, cluster 9 (2,595 genes) and cluster 10 (3,404 genes) start expression at blastulation and remain high afterwards, and are enriched for translation related processes. Meanwhile, clusters 11 (2,708 genes) and 12 (1,911 genes), were primarily associated with somitogenesis and later development, showing enrichment for organ development. Finally, genes with relatively stable expression across all stages, a hallmark of HKGs, were grouped in cluster 8 (2,471 genes) and were enriched for the transcription, translation and cell division related functions.

### Identification of Pluripotency Genes in Atlantic Salmon

Maintenance of pluripotent cells is tightly linked to embryonic development, and the expression of pluripotency maintenance genes progressively decreases during differentiation. We therefore chose to use our dataset to identify genes that control or maintain pluripotency in Atlantic salmon. As a first step we compiled a reference list of known pluripotency factors identified in human, mouse (42,43) and teleosts (50), yielding 286 unique gene names. Due to incomplete annotation of the Atlantic salmon genome, we mapped orthologs across human, zebrafish, and rainbow trout using Ensembl biomaRt, identifying putative orthologs for 240 genes. This resulted in 606 annotated Atlantic salmon gene entries, reflecting extensive salmonid-specific duplication; for instance, orthologues of the gene *klf17* matched to 17 different salmonid genes. Of the 606 genes, 507 genes were expressed in our RNA-seq dataset.

We examined expression patterns of these 507 genes using SOM clustering (Figure 9). A substantial number of genes (260 in clusters 1,2,5,6, and 7) were highly expressed after gastrulation, which precludes their function as regulators or maintainers of pluripotency. In contrast, genes in clusters 10, 11 and 12 showed high expression during pre-gastrulation stages, consistent with a role in maintaining pluripotency. These included canonical pluripotency markers *pou5f3*, *nanog*, *myc*, *sox19a*, *sox19b*, *klf2*, and *klf17* (Supplementary Table 8)(61,62).

**Figure 9.**
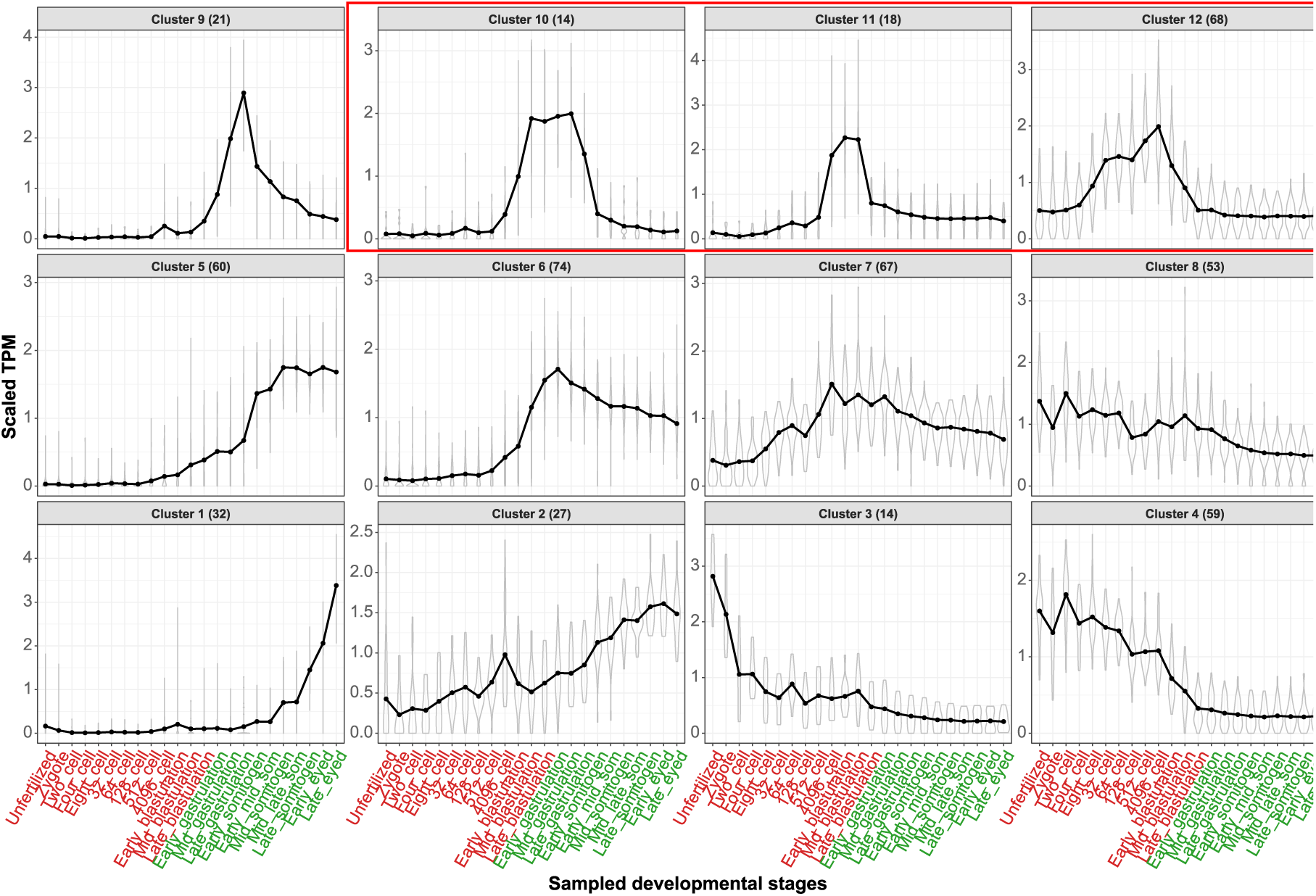
SOM clustering of the potential stem cell markers. Clusters in the red box shows expected transcriptomic profile of pluripotency genes

Considering potential limitations of this candidate gene approach, we also conducted a de novo DE analysis. As the first step in this analysis, we compared gene expression levels between stages associated with pluripotency (4096-cell to late blastula) and those indicative of differentiation (late gastrula to late somitogenesis Of the 54,464 genes analysed, 6,354 (12%) showed significantly higher transcript levels in pluripotency-associated stages (adjusted *p* < 0.05, log₂ fold change > 1), while 10,576 genes were downregulated. GO enrichment analysis of upregulated genes revealed enrichment for functions associated with mRNA 3’-UTR binding, cell cycle regulation, and meiotic nuclear division (Supplementary Table 9), consistent with mechanisms known to sustain undifferentiated states. Notably, 52 of these genes overlapped with our curated list of pluripotency factors, including *pou5f3*, *nanog*, *sox19a*, *sox19b*, *klf2a*, and *myc* (Supplementary Table 8).

Since the mechanisms of pluripotency maintenance are evolutionarily conserved (63), we further refined our candidate gene lists by integrating transcription factor motif analysis with available chromatin accessibility data. Pou5f3/1 is a transcription factor that plays a central role in maintaining pluripotency across vertebrates, binding characteristic motifs in the promoter region of a wide variety of genes to either activate their expression or repress them to maintain the undifferentiated state of the cells. We therefore searched for its conserved DNA binding motif (64) within 2.5 kb upstream and 1 kb downstream of the transcription start site (TSS) across the genome. The range was chosen because Pou5f3/1 is an auto regulator, and its binding sites are generally found withing this 3.5 kb region. For our analysis we chose TSS based on the longest transcript per gene. As a result, we identified 31,635 genes with at least one putative Pou5f3/1 motif within this defined promoter region.

Since the pluripotency factors must be expressed at the stages that contain predominantly pluripotent cells, we used our transcriptomic data to narrow the list to 3,510 genes. Recognizing that this number is still too large to represent core regulators alone, we used the late blastulation stage chromatin state data from the AQUA-FAANG project to filter for motif sites overlapping regulatory regions designated as "Accessible Chromatin," "Active TSS," "Upstream Flanking," or "Active Enhancer" resulting in a set of 7,588 genes with motifs in accessible regions. 1,056 of which were upregulated during pluripotent stages and have motifs in accessible regions. These included *pou5f3*, *nanog*, *sox19b*, and *myc* but not *klf* family members. Motif positions and associated chromatin states are provided in Supplementary Table 10, with selected examples shown in Figure 10.

**Figure 10.**
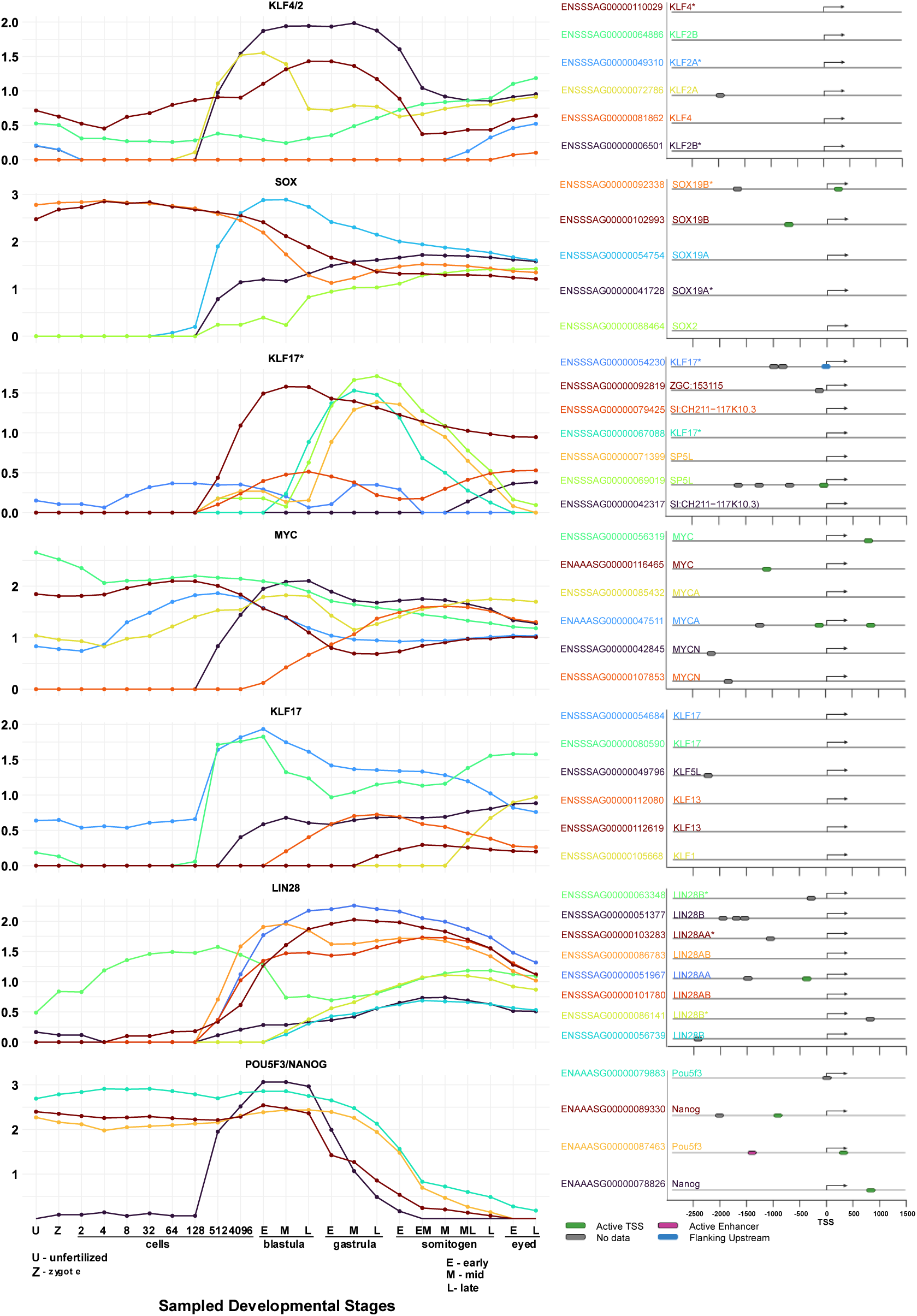
Expression profile of the iPSC related genes and the location of Pou5f3 binding motif in respect to their TSS.

Because Pou5f3/1 often acts in conjunction with Sox family transcription factors, we also searched for genes containing both Pou5f3/1 and Sox2 binding motifs within their promoter regions. We identified 808 such genes, 99 of which were upregulated in the pluripotency stages. Of the 808, only 150 had motifs located in accessible chromatin states, and just 18 were both upregulated and located in regions likely to be transcriptionally active, making them strong candidates for co-regulated pluripotency genes.

### Validation of pluripotency gene expression and HKG identification

To validate gene expression patterns, we chose to examine the levels of the putative pluripotency gene transcripts using qPCR. It is known that for many widely used HKGs, some degree of variation across tissues and experimental conditions exist (65,66). Indeed, levels of the 13 previously reported HKGs (67,68) varied substantially across our samples (Supplementary Figure 6), with only *ub2l3* (ENSSSAG00000092486) showing relatively consistent expression. However, its variance (0.14) was still higher than desired. We therefore proceeded to create a new set of HKG suitable for normalizing gene expression across samples from distinct developmental stages.

We used our RNA-seq data to identify genes with minimal expression variance during the transition from pluripotency to differentiation. Based on scaled read count variance up to early somitogenesis, the gene encoding Ribosomal Operon-Associated A protein (*roaa*) was selected due to its low variance and consistently high expression across early developmental stages (see Supplementary Table 11).

Using *roaa* as a control, we performed qPCR quantification of two pluripotency genes (*nanog*, *pou5f3*) and two potential differentiation markers (*k2o3*, *apoa1*) at late blastulation and early/mid-gastrulation stages. Expression patterns were consistent with RNA-seq results, confirming upregulation of pluripotency markers during blastulation and increased expression of differentiation markers during gastrulation (Figure 11). This observation validates both our RNA-seq based predictions of transcript levels and our HKG normalization strategy for quantifying gene expression in early salmonid embryogenesis.

**Figure 11.**
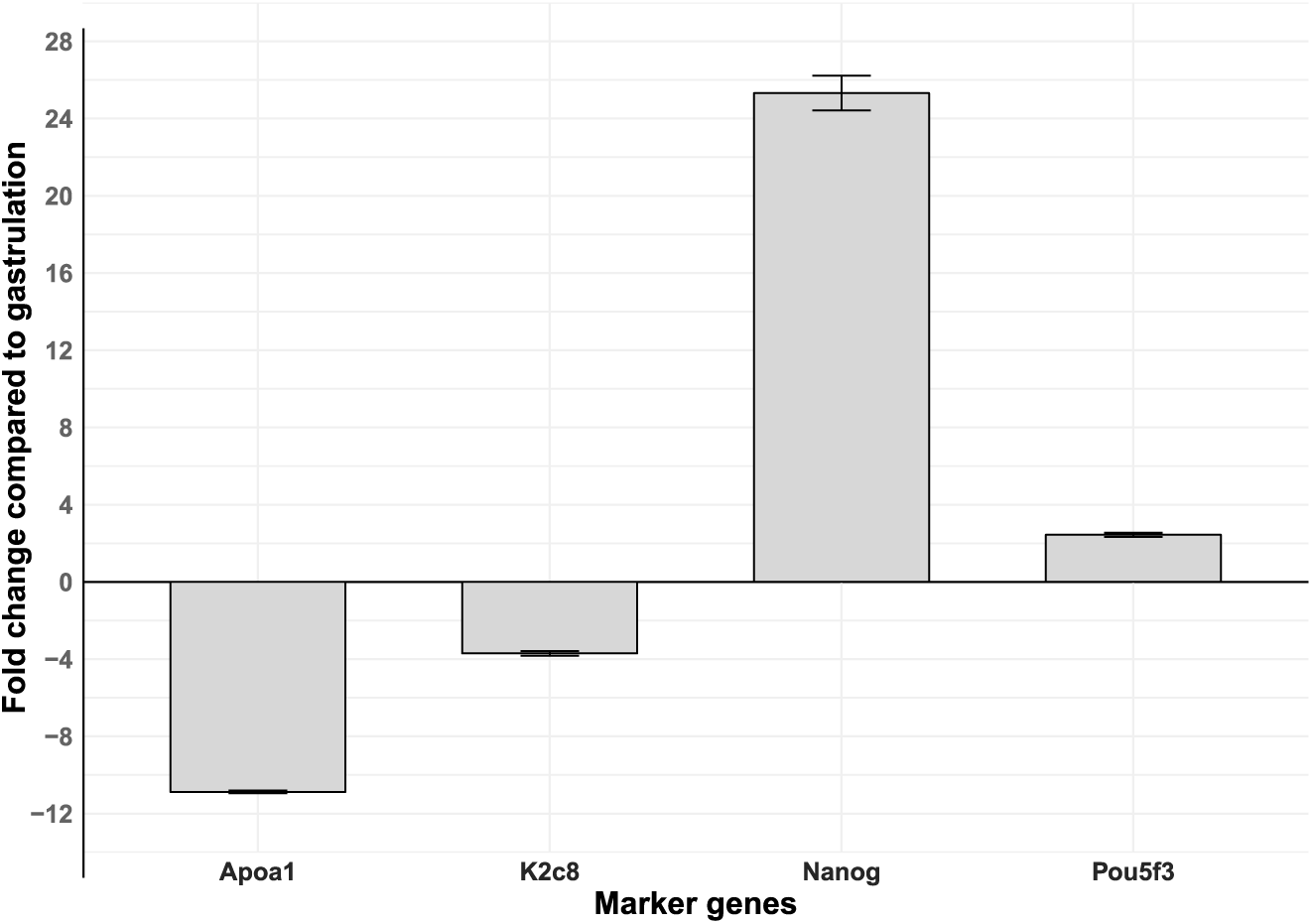
Bar graph of the relative expression of the four marker genes relative to the HKG (roaa) in blastulation compared with gastrulation. Differentiation markers, apoa1 and k2c8, showed downregulation during blastulation compared to the gastrulation. In contrast, pluripotency factors, nanog and pou5f3, were upregulated in the blastulation stage. Standard deviations are indicated by error bars.

Recognizing the need for reliable HKGs, we then calculated the variance in expression for all genes using scaled TPM values across biological replicates (Supplementary Table 12). We selected those with variance < 0.05 (because the scaled TPM ranged between 0 and 4, a variance cutoff of 0.05 corresponds to a standard deviation of only 0.22, or the expression fluctuation by <6% of the full range, indicating highly stable expression across developmental stages), yielding 55 candidate HKGs, all within SOM cluster 7 and 8 (see Figure 8). After removing genes with close homologs (to avoid amplification ambiguity), we identified 7 strong candidates (Table 2). Among them, *H3 histone family 3A (h3f3a)* (ENSSSAG00000095049) had the lowest variance (0.024), which makes it a promising candidate HKG for qPCR normalization of gene expression across Atlantic salmon early developmental stages.

**Table 2.** Best candidate HKGs across Atlantic Salmon embryonic development based on the minimal variance in their expression.

| Gene.stable.ID | Gene description | Gene name | Average TPM | Variance of scaled TPM |
| --- | --- | --- | --- | --- |
| ENSSSAG00000095049 | H3 histone family 3A | *h3f3a* | 160.11 | 0.024 |
| ENSSSAG00000080426 | Ras homolog family member A | *rhoa* | 81.22 | 0.026 |
| ENSSSAG00000012504 | Mitochondrial ribosomal protein S10 | *mrps10* | 38.44 | 0.038 |
| ENSSSAG00000043996 | Translocase of outer mitochondrial membrane 7 | *tom7* | 53.51 | 0.043 |
| ENSSSAG00000022660 | RNA transcription, translation and transport factor | *cnh166* | 85.49 | 0.044 |
| ENSSSAG00000060759 | K1143 protein | *k1143* | 69.85 | 0.045 |
| ENSSSAG00000043062 | PAXIP1 associated glutamate-rich protein 1 | *pagr1* | 25.55 | 0.047 |

## Discussion

Atlantic salmon has recently begun to be extensively studied due to its ecological importance, evolutionary history, and economic value in aquaculture. Despite this, gene expression dynamics during early embryogenesis—particularly the timing of ZGA and the regulation of pluripotency—remain poorly characterized. We chose to close this gap primarily due to our interest in establishing a core set of molecular tools to enable studies of pluripotency in Atlantic salmon. By integrating stage-specific transcriptomic data with orthology-based annotation, differential expression analyses, transcription factor motif scanning, and chromatin state information, we were able to delineate the timing of transition from maternal to zygotic control and identify candidate regulators of pluripotency. Our approach revealed both conserved and novel factors involved in maintenance of pluripotency, while also highlighting the challenges of working with a species with imperfect annotation of a duplicated genome.

To obtain a comprehensive dataset representing a spectrum of early developmental stages we combined information generated in different labs using dissimilar protocols. Evaluation of protocol-specific effects revealed limited variability, with only a small fraction of genes showing differential expression between stranded and unstranded datasets. Consistent with previous reports (41), most differentially expressed genes were overlapping genes on opposite DNA strands. While this supports earlier findings that unstranded library preparation can compromise quantification of overlapping genes, the limited scope of this effect did not impact the integration of these datasets.

### Transitions During Early Atlantic Salmon Development

Our analysis of changes in gene expression across examined developmental stages revealed several general trends. We found concurrent enrichment of cell-cycle regulatory processes and RNA degradation pathways reflects maternal control over early divisions and the progressive clearance of maternal transcripts, a prerequisite for the transition to zygotic regulation (69). At later stages we saw upregulation of genes controlling signaling pathways that establish cell polarity, a key process in early differentiation and tissue morphogenesis (70), and coincided with enrichment of genes involved in organ development (Supplementary Figure 1). Post-gastrulation stages had gene signatures consistent with tissue specialization and rising metabolic demand (18). Notably, the early appearance of genes encoding oxygen-binding components, detectable as early as the 4096-cell stage, identifies a point of a shift toward oxidative metabolism, paralleling observations in mammalian embryos (71).

Together, these findings support a model of coordinated, stage-specific transcriptional programs that guide early embryonic transitions from maternal control toward differentiation and metabolic activation.

### Temporal Gene Clusters Reveal Key Transcriptional Programs in Early Salmon Development

Our analysis of temporal gene expression via unsupervised clustering revealed distinct gene groups associated with maternal, zygotic, and transitional stages. Maternal gene clusters, characterized by high transcript abundance in unfertilized eggs followed by rapid or gradual degradation, were enriched for functions related to cell cycle regulation and DNA polymerase activity. These processes are essential for early embryonic stability and are conserved across vertebrates, including mammals (72). Notably, members of the Smaug (SMG) gene family — regulators of maternal RNA decay (73), were present in these clusters, supporting maternal control of mRNA turnover. Transient expression of complement system components, including complement component 1s, suggests a conserved role for early immune priming in unfertilized eggs, as reported in other teleosts (74,75). Further, the enrichment for function involving the fructose and fatty acid metabolism suggest adaptation towards utilization of yolk.

In contrast, early zygotic gene clusters exhibited enrichment for functions consistent with a transition to embryonic control of gene regulation. A prominent cluster (cluster 6) activated at the four-cell stage (Figure 8) was notably enriched in genes previously observed during the pre–mid-blastula transition in zebrafish (76). At these stages, induction of genes involved in glycerophospholipid metabolism suggests roles in membrane biosynthesis, energy storage, and signalling, reflecting preparatory steps for transcriptional activation and cell division, similar to that observed in mammalian systems (77). During blastulation, appearance of genes related to nutrient response and small nucleolar RNA (snoRNA) activity is consistent with rapid proliferation and preparation for differentiation-associated pathways, including brain and vasculature development during early somitogenesis.

One consideration in interpreting temporal TPM profiles across early development is the effect of global transcriptome remodeling during the MZT. As maternal transcripts are degraded, the total RNA pool contracts, which can cause TPM values of stable or newly transcribed zygotic transcripts to appear elevated not through increased transcription but through reduced competition within the RNA pool. This compositional effect is an inherent property of TPM in the absence of spike-in normalisation and should be considered when interpreting expression dynamics around ZGA. Nevertheless, the coordinated temporal shifts observed across gene clusters align with established features of the maternal-to-zygotic transition, reinforcing that these patterns reflect underlying biological processes rather than purely technical artefacts.

Together, these clustering patterns highlight conserved transcriptional modules underlying the MZT and early embryonic progression, emphasizing the evolutionary stability of core regulatory and metabolic processes across vertebrates.

### Chromatin Remodelling Drives Early and Multi-Phase Zygotic Genome Activation in Atlantic Salmon

ZGA marks a pivotal developmental transition from maternal to embryonic control of gene expression, and its timing varies widely across species (7). One proposed determinant of ZGA timing is the rate of maternal mRNA degradation. In Atlantic salmon, our data reveal that zygotic gene expression begins immediately following fertilization, with additional stage-specific activation of new groups of transcripts as development progresses. This observation supports maternal RNA clearance as a contributing factor to ZGA timing, particularly given that the first cleavage in Atlantic salmon embryos can occur 12-24 hours post-fertilization depending on temperature (78).

The temporal expression profile revealed a sharp increase in zygotic transcription beginning at the zygote stage and peaking during early blastulation. Among the earliest zygotic transcripts were *elp4* and *elp6*, components of the Elongator complex involved in tRNA modification. These genes support translational fidelity during rapid cell proliferation, a conserved function important for both general growth and neurodevelopment (72,73). Early concurrent expression of histone-lysine N-methyltransferases (e.g., *kmt5b* (81), *ehmt1*, and *set1b*) together with transcriptional co-activators such as Cbp (CREB-binding protein), suggests active epigenetic reprogramming via H3K4 methylation, supporting transcriptional activation at the chromatin level (75,76).

Intriguingly, our data suggest a “minor wave” of ZGA occurring as early as the 2- to 8-cell stages, marked by activation of genes with protein kinase activity such as ribosomal protein S6 kinase alpha-3-like (Rps6ka3/Rsk2), which links signalling pathways to transcription via factors including Creb1 (84,85). These stages also saw significant enrichment in genes with histone methyltransferase and deacetylase activity both of which contribute to chromatin remodelling and maintenance of pluripotency (86–88), as seen in early ZGA across vertebrates.

Collectively, our results support a model in which ZGA in Atlantic salmon is both early and multi-phased, with permissive chromatin states and maternal transcript clearance enabling progressive activation of transcriptional programs that underpin pluripotency, cellular homeostasis, and the transition to embryonic control.

### Asynchronous Activation of Ohnologs Highlights Regulatory Divergence During Zygotic Genome Activation

The relatively recent duplication of the salmonid genome provides an opportunity to examine how ohnologs are regulated during early embryogenesis (23). To explore this, we analysed 8,486 genes whose transcripts were initially detected before the early blastula stage and after fertilization. We found that many ohnologs were expressed during this stage, often with substantial asynchrony in the timing of expression within expressed pairs. This asynchrony suggests regulatory divergence or functional specialization of duplicated genes during ZGA. Notably, many maternal genes had ohnolog partners activated zygotically, implying complementary regulation between maternal and zygotic gene pools with similar functional enrichment observed among them.

These findings add to growing evidence that WGD has introduced considerable regulatory complexity in salmonids (89). Divergent temporal regulation of ohnologs during ZGA may represent an evolutionary strategy for fine-tuning gene dosage and function during critical early developmental transitions.

### Evidence for Chromosomal Enrichment During Early Zygotic Activation

Although early embryogenesis is largely governed by maternally deposited factors, increasing evidence suggests that specific chromosomal regions become transcriptionally active during the MZT. In Atlantic salmon, we observed a non-uniform genomic distribution of newly activated zygotic genes, with chromosome 2 consistently overrepresented across multiple early developmental stages. This pattern suggests that certain chromosomes may act as preferential sites for early zygotic activation, paralleling observations in zebrafish (58). It is also an interesting parallel that the chromosomes with these potential MZT gene blocks (chromosome 4 in zebrafish and chromosome 2 in Atlantic salmon) are both sex chromosomes.

Such chromosomal bias may reflect underlying genomic or epigenomic features, including chromatin accessibility, regulatory element clustering, or higher-order genome organization. In zebrafish, early-activated promoters are associated with “placeholder” nucleosomes marked by H2A.Z and hypomethylated DNA, which prime genes for transcription independently of prior activity (90). Comparable mechanisms may operate during the salmon MZT, although direct evidence will require targeted epigenomic and chromatin-conformation analyses along with chromatin accessibility information of earlier developmental stages which is not yet available.

### Dissecting the Pluripotency Architecture in a Genome-Duplicated Vertebrate, Atlantic Salmon

Given that early embryos contain pluripotent cells whose maintenance is tightly linked to embryonic development, identifying the genes that regulate or maintain this state is critical for understanding developmental transitions. As seen in other organisms, early embryonic cells possess totipotency or pluripotency—the ability to give rise to all cell types. This potential is progressively lost as the embryo undergoes gastrulation and cells commit to the ectoderm, mesoderm, or endoderm lineages. Consequently, the onset of gastrulation marks a key transition from pluripotency to lineage restriction.

Understanding pluripotency regulation in non-model vertebrates is also essential for extending stem cell technologies beyond established mammalian systems. In teleosts, iPSC remain largely confined to zebrafish (91), reflecting limited knowledge of species-specific pluripotency networks. In Atlantic salmon, this challenge is compounded by whole-genome duplication, which has expanded and diversified regulatory gene families.

Our analysis of canonical pluripotency-associated genes revealed both conserved and divergent expression patterns compared to mammalian and zebrafish models. While core regulators such as *nanog*, *pou5f3*, *sox19b*, and *lin28ab* exhibited expression profiles consistent with their pluripotency roles, a broader panel of putative Atlantic salmon stem cell markers did not behave as expected. These differences likely reflect both lineage-specific regulatory evolution and constraints of extrapolating pluripotency markers across distant taxa, particularly given that many markers were originally defined in human iPSCs (92), and the regulatory mechanisms can diverge even between model vertebrates like zebrafish and mouse (93,94).

We used the fact - Pou5f3, a conserved master regulator of vertebrate pluripotency (95), to refine a list of candidate pluripotency regulators by identifying genes upregulated during pre- to late-blastulation that contained Pou5f3 binding motifs in their promoter region and then integrating chromatin accessibility data. Our approach resulted in a set that contained known pluripotency genes - *pou5f3*, *nanog*, *sox19b* and *myc*, but notably excludes members of the *klf* gene family.

Given the importance of cooperative transcription factor binding, we further searched for presence of composite Pou5f3/Sox motifs, which yielded a smaller set of high-confidence candidates. The resulting rather small number of salmon genes meeting all our pluripotency criteria, as compared to model organisms (96) likely reflects incomplete chromatin accessibility data at early cleavage stages rather than absence of regulatory interactions. Profiling of earlier stages, such as via ATAC-seq during cleavage, will be necessary to define the full regulatory cascade initiated by Pou5f3 and associated factors.

For our analysis, an additional layer of complexity was presented by the salmonid WGD, that resulted in many genes existing as ohnolog pairs in expanded gene families. These additional gene family members often exhibit non-synchronous expression or display regulatory divergence, complicating interpretation. Although both *nanog* homeologs were expressed during pluripotent stages, only one (*ENSSSAG00000078826*) was associated with accessible Pou5f3/Sox composite motifs. This suggests that the regulation of the *nanog* pair has diverged and one copy is now relying on a different factor to be activated during embryonic development. Similarly, both *pou5f3* homeologs were maternally present but differed in their motif positioning. *sox19b*, rather than the gene currently annotated as *sox2*, emerged as the likely pluripotency-associated *soxB1* factor, mirroring findings in zebrafish. Notably, *ENSSSAG00000081862* is annotated as *klf4*, a key pluripotency factor in mammals, showed little evidence of pluripotency-associated regulation, whereas salmon *klf2a* (*ENSSSAG00000072786*) displayed strong expression and motif support, suggesting possible functional reassignment within the *klf* gene family or misnaming of the specific gene.

Together, these findings reveal a pluripotency architecture in Atlantic salmon that is partially conserved yet extensively reshaped by genome duplication and lineage-specific evolution. While core regulators retain conserved roles, regulatory divergence among duplicated genes underscores the need for functional validation using targeted perturbation and reporter-based assays.

## Conclusions

In summary, this comprehensive temporal transcriptomic analysis of Atlantic salmon embryogenesis provides insights into maternal mRNA clearance, the timing and complexity of zygotic genome activation, and the regulation of pluripotency. By integrating gene expression dynamics with chromatin accessibility and transcription factor motif analyses, we identified conserved and potentially novel pluripotency-associated regulators. This network, however, is shaped by the added regulatory complexity of WGD in salmonids, which introduces expression divergence and functional specialization among ohnologs.

Despite these advances, key questions remain, including the identity of core pluripotency regulators, the regulatory architecture governing early ZGA, and the extent of functional divergence among duplicated genes. These challenges reflect both biological complexity and current technical limitations, including incomplete chromatin accessibility data at pre-ZGA stages and sparse functional annotation in Atlantic salmon.

Future studies leveraging single-cell RNA-seq and ATAC-seq during cleavage and early blastulation stages, as demonstrated in zebrafish (97,98) will be essential for resolving cell-type-specific transcriptional programs and identifying the earliest acting regulatory factors. Moreover, functional validation using CRISPR/Cas9-mediated gene perturbation in salmon embryos, along with enhancer-reporter assays to test motif activity in vivo, will be crucial to move from correlation to causation. Collectively, our work provides a foundational resource for studying vertebrate pluripotency and highlights the importance of resolving regulatory and evolutionary complexity in non-model organisms.

## Acknowledgements and Funding

This work was supported by the LiceResist project, funded by the Research Council of Norway under the HAVBRUK2 programme (project no. 301685), and by the AQUA-FAANG project, which received funding from the European Union’s Horizon 2020 Research and Innovation Programme under Grant Agreement No. 817923 (www.aqua-faang.eu). The authors also gratefully acknowledge internal infrastructure funding provided by the Norwegian University of Life Sciences (NMBU).

