## Supplementary figures 1 - 6 for "Transcriptomic view of key events during early embryogenesis in the duplicated Atlantic salmon genome"

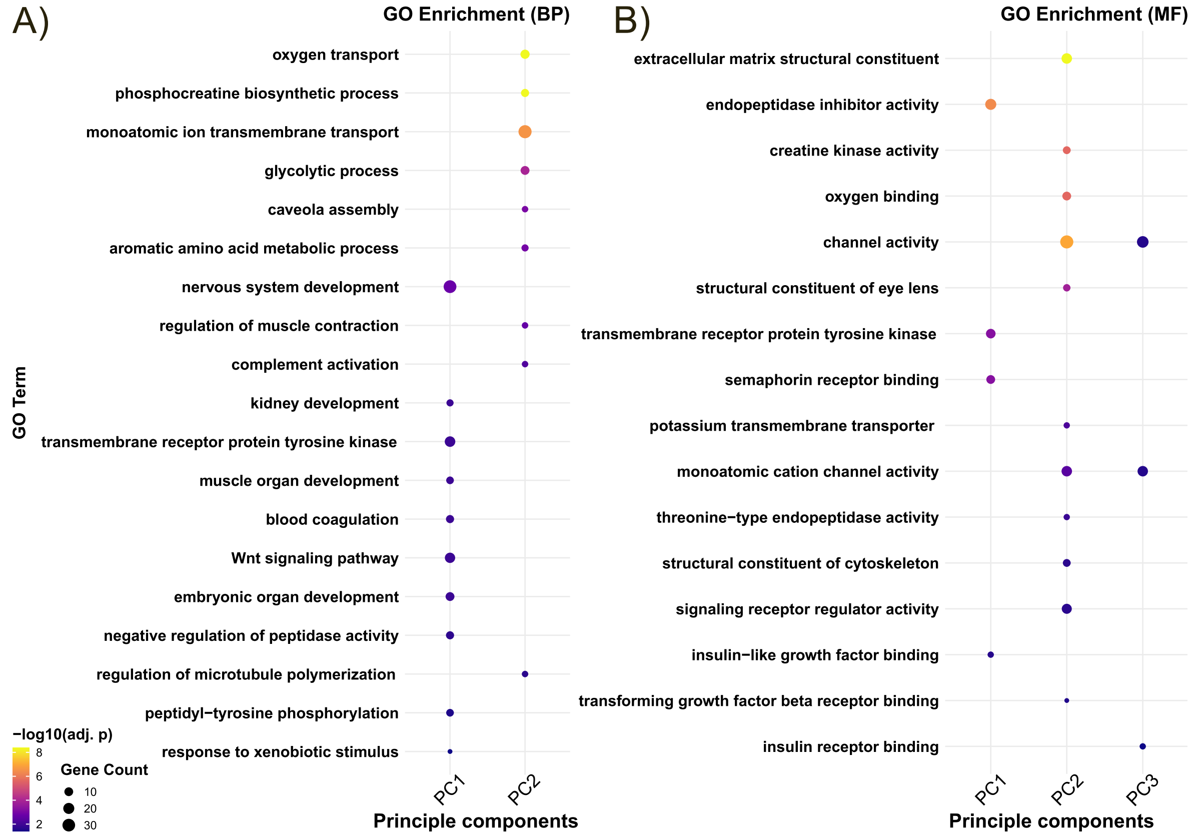


***Supplementary Figure 1*** *Gene Ontology analysis of genes with highest or lowest loading values in the first 3 principal components. A) Biological Process term enriched. B) Molecular Functions enriched in the selected set of genes.*

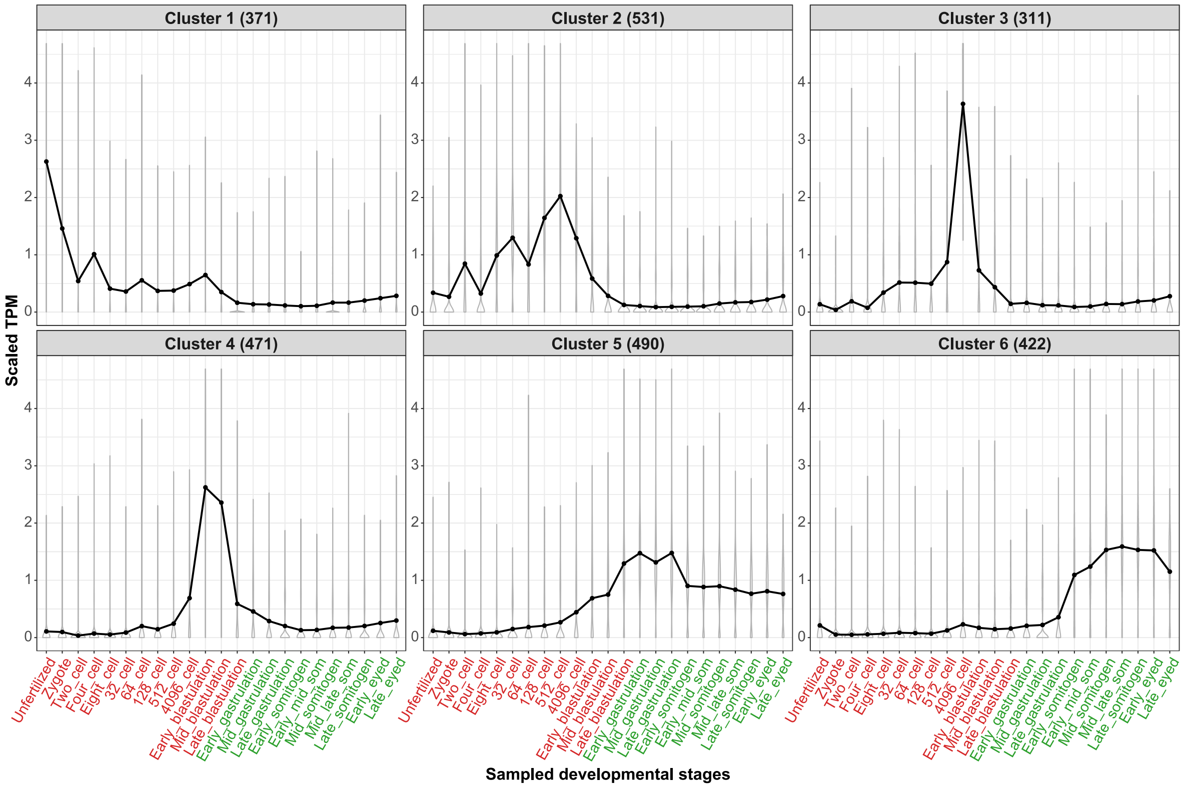


***Supplementary Figure 2*** *SOM clustering of transiently expressed genes (TPM>2). The line goes through the mean scaled TPM for each stage in each developmental stage.*

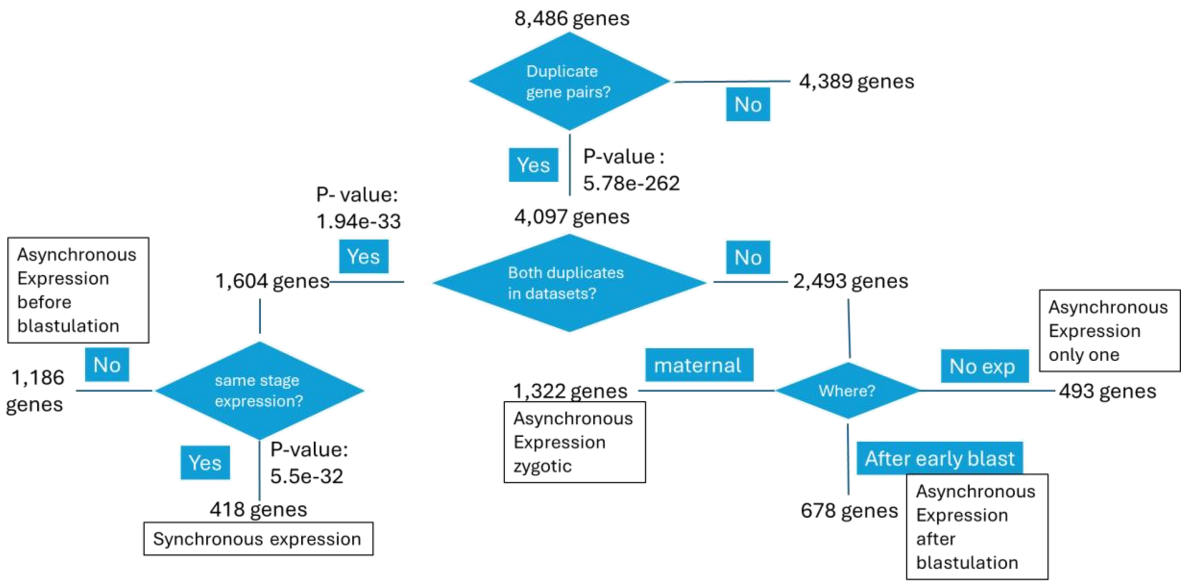


***Supplementary Figure 3*** *Flowchart summarizing the gene expression dynamics for ohnologs among the early expressed genes.*

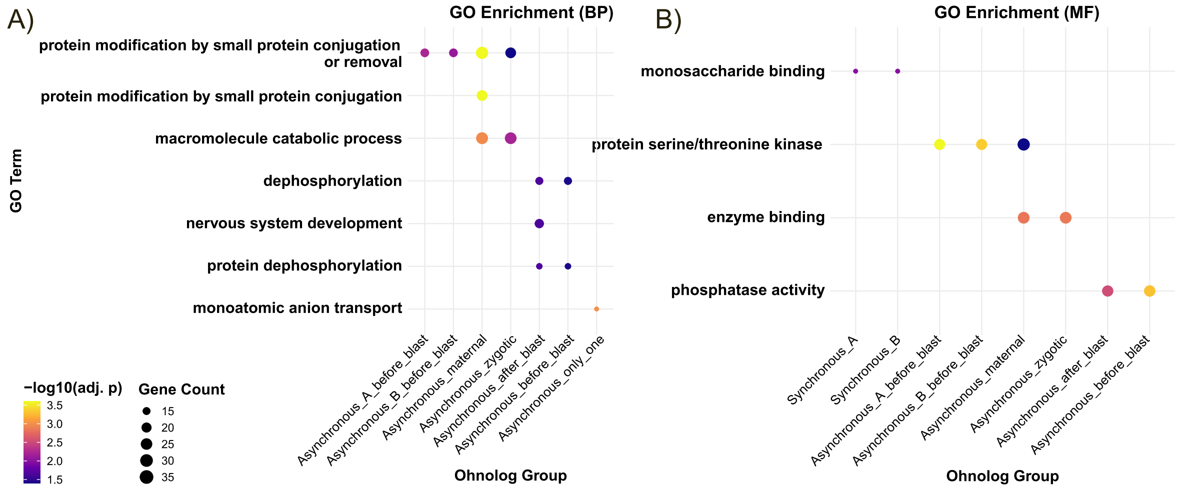


***Supplementary Figure 4 GO analysis of different groups of ohnolog genes expressed during early embryogenesis****. The ohnolog groups are: Asynchronous_A_before_blast- gene set of one of the copy of the ohnolog pair that starts expression before blastulation , Asynchronous_B_before_blast – gene set with the complementary copy of the Asynchronous_A_before_blast, Asynchronous_maternal – gene set of the ohnolog pair whose transcript were detected in unfertilized egg, Asynchronous_zygotic – complementary gene set to Asynchronous_maternal that show expression before blastulation , Asynchronous_after_blast - gene set of ohnologs pair that show expression after blastulation, Asynchronous_before_blast – complementary gene set to Asynchronous_after_blast, Asynchronous_only one – ohnologs where only one in the pair of genes is expressed.*

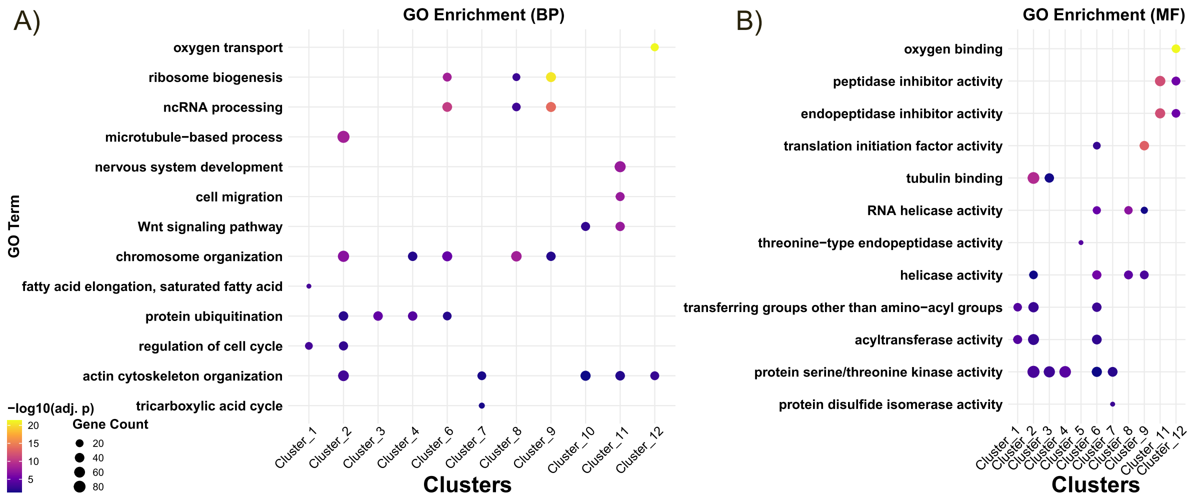


***Supplementary Figure 5*** *Gene ontology enrichment of different clusters of genes. A) Biological processes enriched. B) Molecular functions enriched.*

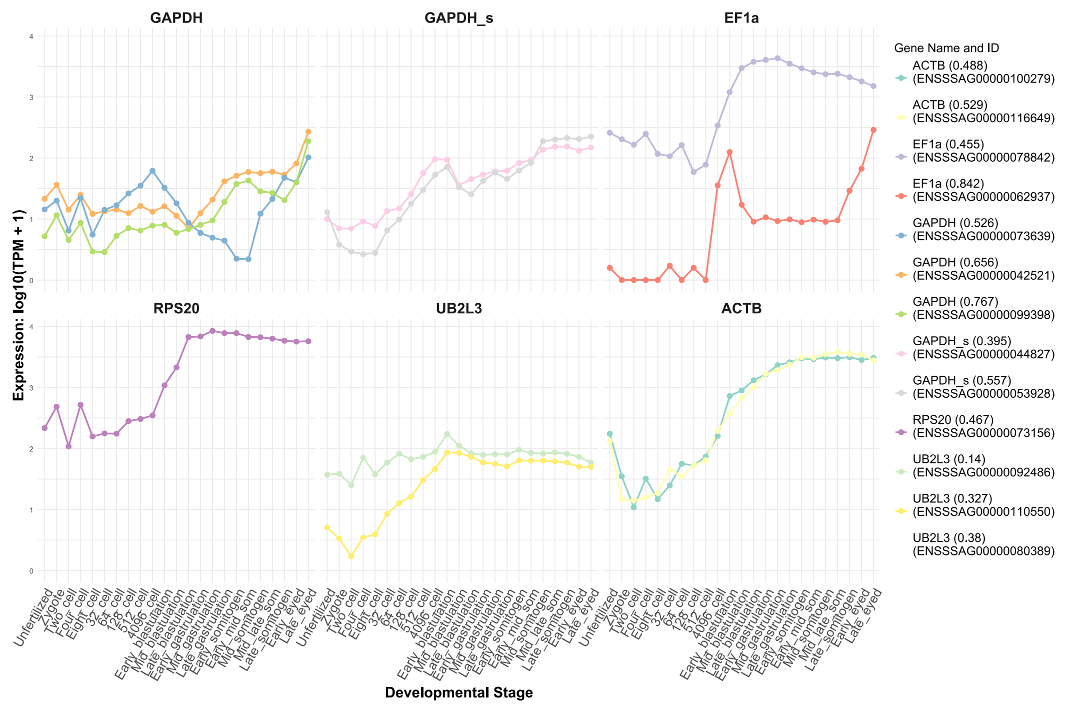


***Supplementary Figure 6*** *Expression profile of previously reported housekeeping genes through embryogenesis along with the observed variance.*
